# Structural basis for aminoacylation of cellular modified tRNA^Lys3^ by human lysyl-tRNA synthetase

**DOI:** 10.1101/2024.12.07.627298

**Authors:** Swapnil C. Devarkar, Christina R. Budding, Chathuri Pathirage, Arundhati Kavoor, Cassandra Herbert, Patrick A. Limbach, Karin Musier-Forsyth, Yong Xiong

## Abstract

The average eukaryotic tRNA contains 13 posttranscriptional modifications; however, their functional impact is largely unknown. Our understanding of the complex tRNA aminoacylation machinery in metazoans also remains limited. Herein, using a series of high-resolution cryo-electron microscopy (cryo-EM) structures, we provide the mechanistic basis for recognition and aminoacylation of fully-modified cellular tRNA^Lys3^ by human lysyl-tRNA synthetase (h-LysRS). The tRNA^Lys3^ anticodon loop modifications S34 (mcm^5^s^2^U) and R37 (ms^2^t^6^A) play an integral role in recognition by h-LysRS. Modifications in the T-, variable-, and D-loops of tRNA^Lys3^ are critical for ordering the metazoan-specific N-terminal domain of LysRS. The two catalytic steps of tRNA^Lys3^ aminoacylation are structurally ordered; docking of the 3′-CCA end in the active site cannot proceed until the lysyl-adenylate intermediate is formed and the pyrophosphate byproduct is released. Association of the h-LysRS-tRNA^Lys3^ complex with a multi-tRNA synthetase complex-derived peptide shifts the equilibrium towards the 3′-CCA end ‘docked’ conformation and allosterically enhances h-LysRS catalytic efficiency. The insights presented here have broad implications for understanding the role of tRNA modifications in protein synthesis, the human aminoacylation machinery, and the growing catalog of metabolic and neurological diseases linked to it.

## INTRODUCTION

Transfer ribonucleic acids (tRNAs) are essential non-coding adaptor RNA molecules that mediate the fidelity of mRNA translation across all living organisms(1). Amino acids (aa) required for protein synthesis are delivered to the ribosome by aminoacylated or ‘charged’ tRNAs, activated forms of tRNAs wherein the 3’-end is conjugated with a cognate aa by specialized enzymes called aminoacyl-tRNA synthetases (AARS)(2). Almost all organisms encode at least 20 AARS, one for each aa, whereas higher eukaryotes encode a distinct set of mitochondrial and cytosolic AARS(3,4). Lysyl-tRNA synthetase (LysRS) is one of three human (h) AARS, wherein both the mitochondrial and cytosolic isoforms are encoded by the same gene and produced via alternative splicing(4). In most organisms, LysRS belongs to the class IIb subfamily of AARS and is evolutionarily and structurally related to aspartyl-tRNA synthetase (AspRS) and asparaginyl-tRNA synthetase (AsnRS)(5,6). All class II synthetases possess oligomeric structures (dimers or tetramers) and contain three highly conserved active site motifs(7). Motif 1 makes up the dimer interface and motifs 2 and 3 make up the aminoacylation active site, a seven-stranded b-sheet structure. In contrast to its bacterial counterpart, h-LysRS has a 70 aa N-terminal domain (NTD) that facilitates tRNA binding(8) but the structural basis of its role in aminoacylation remains unresolved. In metazoans, almost half of all cytoplasmic AARS and three scaffold proteins called ‘aminoacyl-tRNA synthetase complex interacting multifunctional proteins’ (AIMP) assemble to form the multi-tRNA synthetase complex (MSC) (Figure 1A)(9–11). The exact architecture of the MSC and its regulatory role in aminoacylation remains controversial(11,12). h-LysRS is associated with the MSC via an interaction with AIMP2 (p38) and is released from the MSC after phosphorylation at S207(13,14). pS207-LysRS is a regulator of nuclear transcription through synthesis of the dinucleotide Ap4A and is inactive for tRNA^Lys^ aminoacylation(15,16).

**Figure 1.**
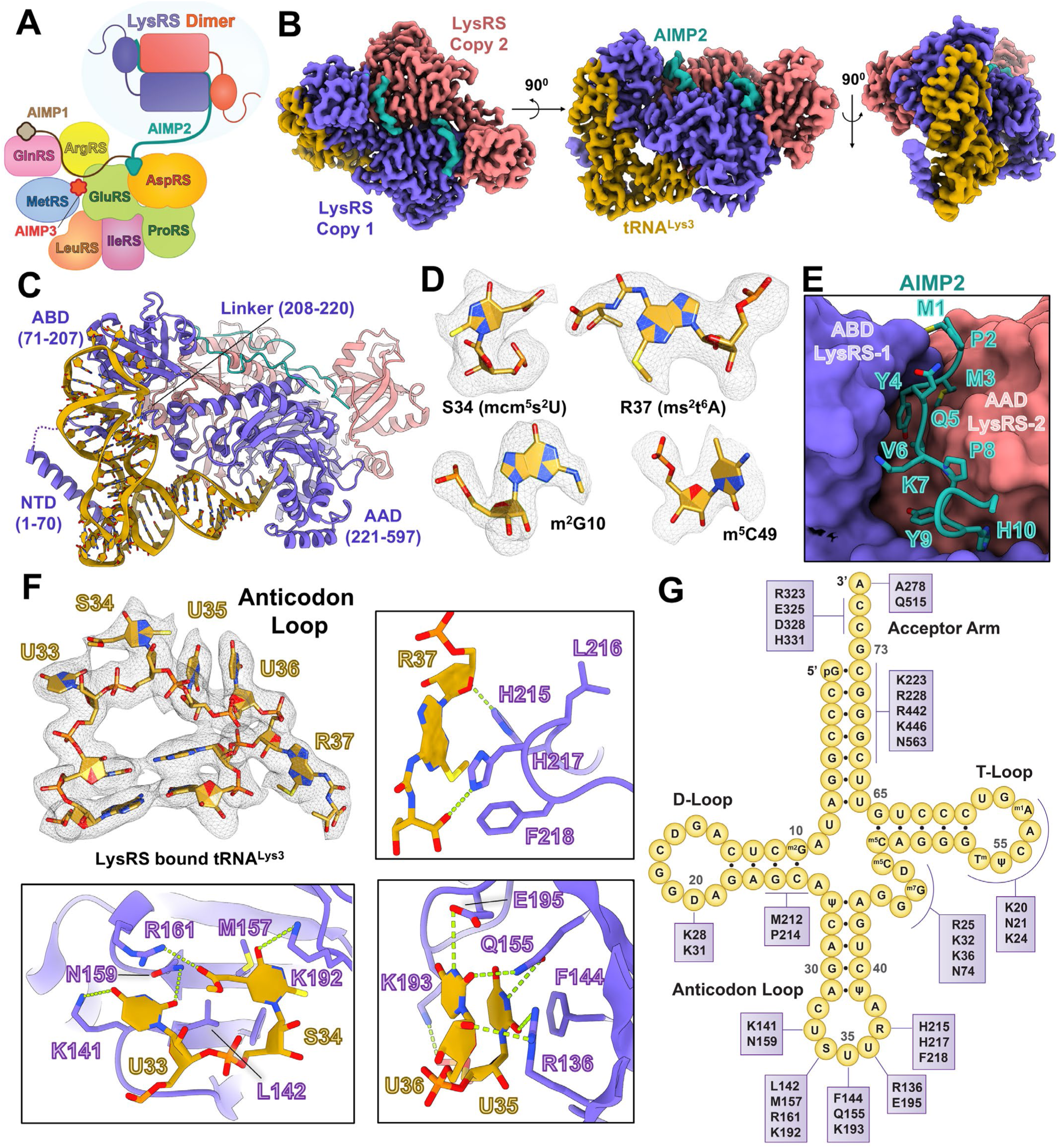
Structure of h-LysRS bound to modified cellular tRNA^Lys3^ and AIMP2. (A) A schematic representation of the multi-tRNA synthetase complex (MSC). h-LysRS dimer is anchored to the MSC by AIMP2 and is highlighted. (B) Cryo-EM reconstruction of wt h-LysRS bound to modified cellular tRNA^Lys3^ and AIMP2 in the presence of AMP and L-Lysine. (C) Structural model built for the h-LysRS-tRNA^Lys3^-AIMP2 complex based on the cryo-EM reconstruction shown in panel B. The various domains of h-LysRS are highlighted (NTD – N-terminal domain; ABD – Anticodon binding domain; AAD – Aminoacylation domain). (D) Representative modified nucleotides in tRNA^Lys3^ are shown with the corresponding cryo-EM map density from the reconstruction shown in panel B. (E) The N-terminal residues of AIMP2 (teal) secure the dimeric interface between the ABD of one LysRS monomer (purple) and the AAD of the other (coral orange) and are shown in stick representation. (F) The anticodon loop of tRNA^Lys3^ (U33-R37) bound to h-LysRS is shown along with the corresponding cryo-EM map density. The insets highlight the interactions of the anticodon loop nucleotides (gold) with LysRS (purple). (G) The interaction map of the h-LysRS-tRNA^Lys3^ complex is shown with tRNA^Lys3^ portrayed in the conventional clover-leaf representation.

Another layer of complexity in higher eukaryotes is the extensive repertoire of RNA modifications observed in tRNAs, with an average of 17% of tRNA bases being modified post-transcriptionally in humans(17,18). tRNA^Lys3^(UUU), a major species of tRNA^Lys^ in humans, carries two RNA modifications in its anticodon loop - 5-methoxycarbonylmethyl-2-thiouridine (mcm^5^s^2^U) on base 34 (S34) and 2-methylthio-N6-threonylcarbamoyladenosine (ms^2^t^6^A) on base 37 (R37)(19). These modifications have been shown to regulate codon-anticodon wobble base-pairing and ribosomal frameshifting(20–22). tRNA^Lys3^ also carries an array of RNA modifications in the D-loop, T-loop, and variable loop regions that govern RNA folding and stability(17,19,23,24). However, the exact role of these modifications in recognition and aminoacylation by h-LysRS remains unknown.

Mutations in the cellular aminoacylation and RNA modification machineries have been linked to a growing number of neurodegenerative and metabolic disorders(4,25–27). h-LysRS has been implicated in Charcot-Marie-Tooth disease(28) and congenital leukoencephalopathy(29), whereas mutations in enzymatic machinery carrying out the ms^2^t^6^A modification on tRNA^Lys3^ are linked to type II diabetes(30) and Galloway-Mowat syndrome(31).

tRNA^Lys3^ also serves as the primer for HIV-1 reverse transcription and the packaging of the h-LysRS-tRNA^Lys3^ complex is an indispensable step in the HIV-1 lifecycle(32). Structural studies on h-LysRS have been limited to crystallization and NMR studies of its apo-form, often using truncated versions of the protein with the NTD missing in all the structures reported to date (6,13,33). The only available structural information on the LysRS-tRNA^Lys^ complex comes from a *Thermus thermophilus* LysRS-tRNA^Lys(UUU)^ complex(34). However, in this structure, only the anticodon stem-loop of bacterial tRNA^Lys^ was resolved in complex with the anticodon binding domain; how LysRS interacts with the acceptor stem and the 3’-CCA end remains unknown. Furthermore, bacterial LysRS lacks the NTD of h-LysRS and tRNA^Lys3^ in humans is more extensively modified compared to its bacterial counterpart. Thus, the structural and mechanistic basis for tRNA^Lys3^ recognition and aminoacylation by h-LysRS remains elusive.

To address these fundamental gaps in our understanding of tRNA^Lys3^ recognition by h-LysRS and the role of tRNA^Lys3^ modifications and the MSC in aminoacylation, we carried out a comprehensive structural and enzymatic characterization of h-LysRS complexed with cellular modified tRNA^Lys3^ and the MSC scaffold protein AIMP2. Our results provide a structural and mechanistic basis for tRNA^Lys3^ specificity of h-LysRS and the conformational landscape of the LysRS catalytic pathway. A comparison of h-LysRS complexes with cellular modified tRNA^Lys3^ and *in vitro* transcribed (IVT) tRNA^Lys3^ (unmodified), revealed how tRNA^Lys3^ modifications are directly involved in recognition by h-LysRS and how the absence of these modifications compromises recognition. Strikingly, the two steps in the h-LysRS aminoacylation reaction are ordered structurally, wherein the 3’-CCA end docking in the active site cannot proceed until the ATP hydrolysis-driven L-lysyl adenylate intermediate is formed and the resulting pyrophosphate moiety is released. AIMP2, which anchors h-LysRS to the MSC, allosterically enhances the catalytic efficiency of h-LysRS by driving the equilibrium of the h-LysRS-tRNA^Lys3^ complex towards the 3’-CCA-end ‘docked’ conformation. The detailed mechanistic and structural insights into tRNA^Lys3^ aminoacylation by h-LysRS and the role of tRNA modifications and the MSC have broad implications for understanding the aminoacylation machinery in humans and the growing number of neurological and metabolic disorders linked to it.

## RESULTS

### Structure of human LysRS-cellular modified tRNA^Lys3^ -AIMP2 complex

To gain a mechanistic and structural understanding of tRNA^Lys3^ recognition and aminoacylation by LysRS, we carried out cryo-EM studies of h-LysRS complexed with cellular modified tRNA^Lys3^ and the MSC adaptor protein, AIMP2. Fully modified tRNA^Lys3^ was purified from HEK293T or Jurkat cells (detailed protocols in Methods section) and mass spectrometry analysis validated the presence of most of the previously reported modifications in the purified tRNA^Lys3^ sample (Supplementary Figure S1A-B and Supplementary Table S1)(19). The LysRS interacting N-terminal region of AIMP2 (N36) was chemically synthesized and used in the cryo-EM studies(13). Cellular modified tRNA^Lys3^ was complexed with recombinantly purified h-LysRS and AIMP2-N36 in the presence of adenosine monophosphate (AMP) and L-lysine for single-particle cryo-EM structural studies. A final reconstruction of dimeric LysRS bound to a single modified tRNA^Lys3^ and two copies of AIMP2-N36 was resolved to 2.75 Å and used for model building (Figure 1B-C, Supplementary Figure S1C-F, Supplementary Table S2-S4).

The high-resolution cryo-EM reconstruction allowed for unambiguous modeling of majority of h-LysRS, AIMP2-N36, and all nucleotides of the cellular modified tRNA^Lys3^ (Figure 1C, Supplementary Figure S2A, and Supplementary Table S4). Consistent with the mass spectrometry analysis, tRNA^Lys3^ modifications exhibited distinct densities in the cryo-EM reconstruction (Figure 1D). The only tRNA^Lys3^ modifications that cannot be confirmed by mass spectrometry analysis are pseudouridines (ψ) due to their identical mass to uridines. However, the H1N1 imino group of pseudouridine is a strong hydrogen bond donor and often leads to a water molecule being coordinated between its 5’-phosphate and H1N1 imino group(35,36). This molecular signature was used to confirm the presence of pseudouridine in the cellular modified tRNA^Lys3^ used in this study (Supplementary Figure S2B). Nine different tRNA modifications and a total of 14 modified nucleotides were modeled for the cellular purified tRNA^Lys3^.

The first ten residues of AIMP2 act as a latch for securing the dimeric interface formed by the anticodon-binding domain (ABD) of one LysRS monomer with the aminoacylation domain (AAD) of the other and anchors the LysRS-tRNA^Lys3^ complex to the MSC (Figure 1E). Strikingly, the anticodon loop (33–37) is extensively remodeled by LysRS, compared to the crystal structure of tRNA^Lys3^ alone (PDB: 1FIR)(37) (Figure 1F and Supplementary Figure S2C). All five bases in the anticodon loop are unstacked and form an extensive network of interactions with LysRS (Figure 1F and 1G). Interestingly, U33 and R37, adjacent to the anticodon bases S34, U35, and U36, are also integral to the recognition of tRNA^Lys3^. U33 forms base-specific salt bridges with K141 and N159 while stacking against the aliphatic sidechain of K141 and the guanidium group of R161. The RNA modifications on S34 (mcm^5^s^2^U) form specific interactions with the LysRS ABD, wherein R161 forms a salt bridge with the 5-mcm moiety and the 2-thiol modification stacks against T191 and the aliphatic sidechain of K193 (Figure 1F). These interactions are further strengthened by hydrophobic packing interactions of S34 against L142, F144, and M157. U35 and U36 are recognized by LysRS ABD via base-specific ionic interactions as well as hydrophobic stacking interactions. F144 forms a three-ring stack with U35 and U36, which is further stabilized by salt bridges between R136, Q155, and E195 of LysRS ABD and U35 and U36 of tRNA^Lys3^, respectively (Figure 1F). The ms^2^t^6^A-modified base (R37) is recognized by the linker region (208–220) connecting the LysRS ABD to its AAD. H217 and F218 of this linker region stack against the adenine base and the bulky N6-threonylcarbomyl modification, whereas H215 forms hydrophobic stacking interaction with the 2-methylthio moiety of R37 (Figure 1F).

The hydrophobic stacking interactions of H215 and H217 are further strengthened by hydrogen bonding with the R37 phosphate backbone and the N6-threonylcarbomyl modification, respectively.

The NTD of LysRS (1–65) is unique to higher eukaryotes and, in the absence of tRNA^Lys3^, is predicted to be unstructured(38). The NTD of h-LysRS is either disordered or was deleted in all the X-ray crystallography structures reported to date(6,13). h-LysRS NTD is highly basic and has been proposed to play a role in tRNA binding and aminoacylation(8). In the complex structure reported here, residues 14-43 of the NTD are clearly resolved in the cryo-EM reconstruction and form an extended α-helix that is anchored at the base of the tRNA^Lys3^ T-loop and runs along the anticodon arm of tRNA^Lys3^ (Figure 2A and Supplementary Figure S2D). All the basic residues of the N-terminal helix face tRNA^Lys3^, creating a highly positive surface that mediates several key protein-RNA interactions (Figure 1G, 2A-B, and Supplementary Figure S2E). Residues K20, at the N-terminus of the extended helix, N21, and K24 interact with the phosphate backbone of the T-loop. K20 and K24 also form ionic interactions with the modified nucleotide ψ55. In addition, K28, K32, and K36 interact with the phosphate backbone of the anticodon arm of tRNA^Lys3^. Interestingly, two key RNA modifications in the D-loop and the variable loop play crucial roles in stabilizing the N-terminal helix. U20 of the D-loop and U47 of the variable loop are modified to 5,6-dihydrouridine (D), leading to a loss in the planar nature of these pyrimidine bases and loss in their ability to stack against the tRNA^Lys3^ central core. In the structure reported here, D20 and D47 of tRNA^Lys3^ adopt an unstacked conformation, creating a wider groove between the D-loop and the variable loop and allowing the NTD of h-LysRS to dock. D20 and D47 further stabilize the h-LysRS NTD by forming interactions with K31 and R25, respectively (Figure 2B).

**Figure 2.**
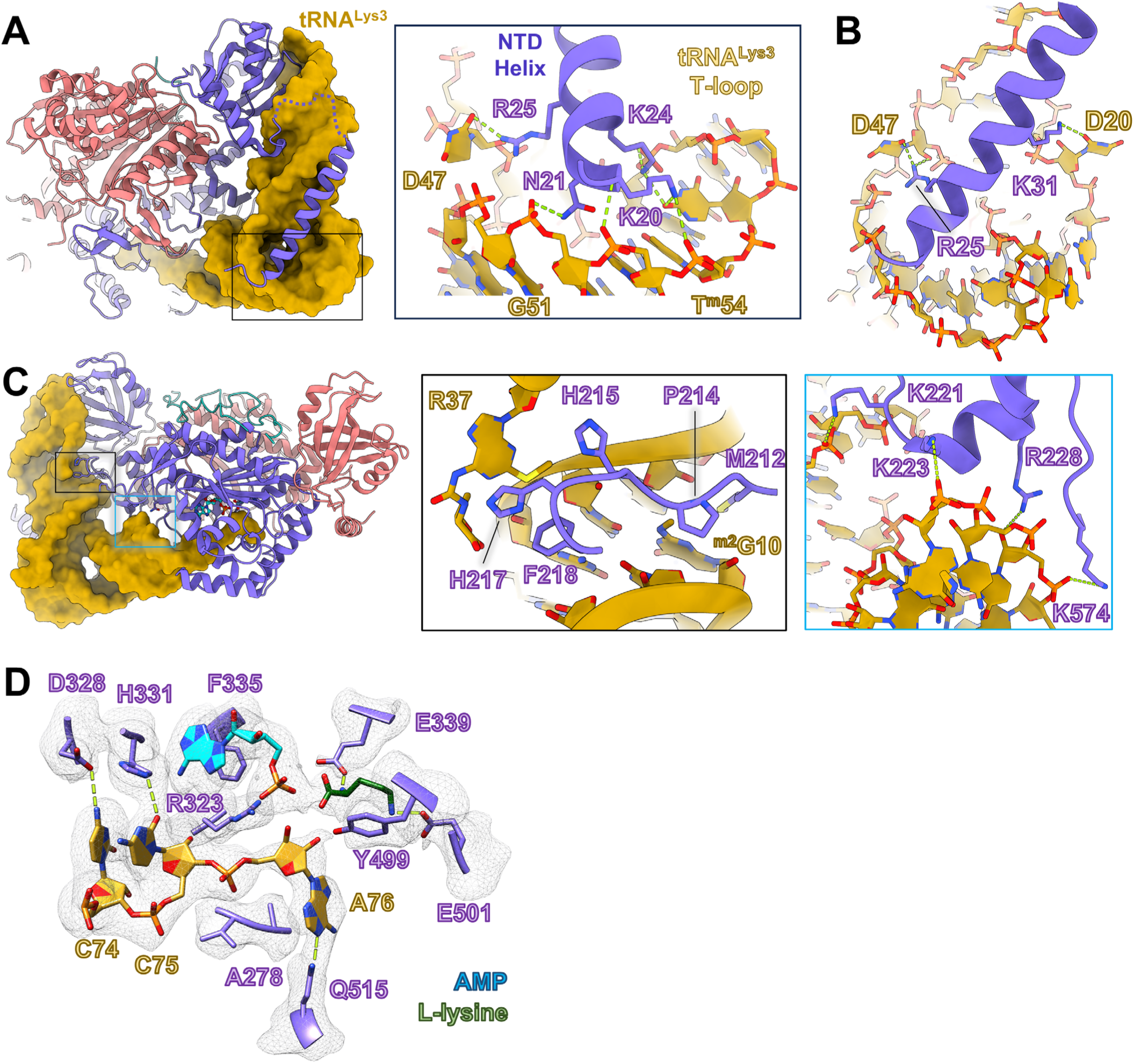
tRNA^Lys3^ interactions with h-LysRS NTD and active site. (A) A view of the h-LysRS-tRNA^Lys3^ complex (h-LysRS in cartoon and tRNA^Lys3^ in surface view) showing the NTD helix of LysRS interacting with the elbow region of tRNA^Lys3^. The inset highlights the residues at the base of the NTD helix and their contacts in the tRNA^Lys3^ T-loop and variable loop. (B) A magnified view showing the N-terminal helix of h-LysRS and its interactions with two dihydrouridine bases of tRNA^Lys3^, D2O and D47, of the D-loop and variable loop, respectively. (C) The docked state h-LysRS-tRNA^Lys3^ complex is shown, with the insets highlighting key interactions for docking the 3’-CCA end of tRNA^Lys3^. LysRS linker residues interact with the D-arm (middle) whereas the acceptor arm is stabilized by a series of ionic interactions (right). (D) A magnified view of the h-LysRS active site is shown with the tRNA^Lys3^ 3’-CCA end docked in. The corresponding cryo-EM map density is shown in mesh.

Similar to all class II AARS, the active site of h-LysRS is formed at the base of six antiparallel β-sheets and a seventh parallel β-sheet. The h-LysRS active site has three pockets, one for each of the substrates in the aminoacylation reaction – ATP, L-lysine, and the 3’-CCA end of the tRNA. The cellular modified tRNA^Lys3^ in this reconstruction was present exclusively in the ‘docked’ state, wherein the 3’-CCA end is inserted into the h-LysRS active site. The key interactions for positioning the acceptor arm towards the active site involve h-LysRS linker residues M212 and P214 that contact tRNA^Lys3^ from the D-arm minor groove and form hydrophobic stacking interactions with 2-methyl guanosine (m^2^G10) as well as F218 and K221 that form additional interactions with the D-arm (Figure 2C). The acceptor arm is stabilized by interactions mediated by K223, E224, R228, R442, K446 and K574 (Figure 2C). All three 3’-terminal residues of the tRNA form base-specific interactions with h-LysRS AAD (Figure 2D). C74 and C75 of tRNA^Lys3^ are recognized by h-LysRS motif 2 residues. The base of C74 forms a hydrogen-bonding network with E325, G326, and D328. In the active site, H331 coordinates C75 by forming base-specific interactions while simultaneously stacking against the adenine base of AMP. E325 and H331 are invariable motif 2 residues of LysRS conserved across prokaryotes and eukaryotes(39). The 3’-terminal A76 base is unstacked relative to C75 and stabilized in a hydrophobic pocked formed by the ‘flipping loop’ (272–282), located between motifs 1 and 2, and residues P471, L472, and F512. The flipped A76 base stacks on A278 and forms hydrogen bonds with Q515. This conformation of A76 places the 3’-hydroxyl group of the ribose near the α-phosphate of AMP and the α-carboxylic group of L-lysine. An alternate conformation for the A76 base (conformation B) was also observed in the cryo-EM reconstruction wherein the A76 base is rotated away from the active site and placed in an alternate hydrophobic pocket formed by A278, V279, F528, and I529 (Supplementary Figure S2F). The functional relevance of this alternate A76 binding pocket in aminoacylation is not apparent but may have a role in Ap4A synthesis carried out by h-LysRS. Both conformations of the flipped A76 base are stabilized by hydrogen bonding between the amino group of A76 base and Q515 of h-LysRS.

The binding mode of ATP and L-lysine in the h-LysRS active site is highly conserved across species as well as within the class IIb family of AARS(6,40). The adenine base of the AMP forms pi-stacking interactions with F335 and is further stabilized by H331 and R553. The class II invariant motif 2 residue R323 coordinates the α-phosphate of AMP and is within hydrogen bonding distance from the 3’-hydroxyl of the A76 ribose. L-lysine is coordinated in the h-LysRS active site via a series of ionic and hydrophobic interactions. The aliphatic sidechain of L-lysine stacks in between Y341 and Y499 whereas the ε-amino group interacts with E501. E339 and E301 form ionic interactions with the α-amino group of L-lysine, placing its α-carboxylic group near the α-phosphate of AMP and the 3’-hydroxyl group of the A76 ribose (Figure 2D).

### ATP state governs 3’-CCA end docking of tRNA^Lys3^ in the active site

To gain a mechanistic understanding of the aminoacylation pathway for tRNA^Lys3^, we carried out cryo-EM studies of the h-LysRS-tRNA^Lys3^ complex in the presence of a non-hydrolyzable ATP analog, α,β-methylene-adenosine 5’-triphosphate (AMPCPP), as well as AMP to mimic the pre- and post-hydrolysis states of LysRS-tRNA^Lys3^ complex, respectively. The tRNA^Lys3^ used in these studies was produced by *in vitro* transcription (IVT), and therefore, lacked any RNA modifications. This allowed us to systematically study the effects of individual cellular tRNA^Lys3^ modifications on the h-LysRS-tRNA^Lys3^ complex. The AMP dataset yielded three distinct reconstructions - apo-LysRS, LysRS-tRNA^Lys3^ complex in the ‘undocked’ state, and h-LysRS-tRNA^Lys3^ complex in the ‘docked’ state (Figure 3A and Supplementary Table S5). In contrast, the conformational state of the LysRS-tRNA^Lys3^ complex in the AMPCPP dataset was exclusively in the ‘undocked’ state (Figure 3A and 3B). Compared to the ‘docked’ state, the LysRS active site in the AMPCPP bound ‘undocked’ state has notable differences. AMPCPP assumes a bent conformation in the LysRS active site, wherein the β- and γ-phosphate fold back towards the adenine base (Figure 3B). The interactions of the adenine base of AMPCPP are similar to that observed in the docked state detailed above. Two magnesium ions were observed in the LysRS active site in the AMPCPP bound ‘undocked’ state, coordinated by conserved residues E487 and E494, and play a key role in orienting the triphosphate moiety of ATP in the LysRS active site. In the presence of AMP, no discernable density attributable to magnesium ions was observed in the h-LysRS active site. This difference between the AMPCPP and AMP bound states of h-LysRS suggests that magnesium plays an active role in the first catalytic step of h-LysRS, ATP hydrolysis and formation of the lysyl-adenylate intermediate, but is not involved in the second step of catalysis.

**Figure 3.**
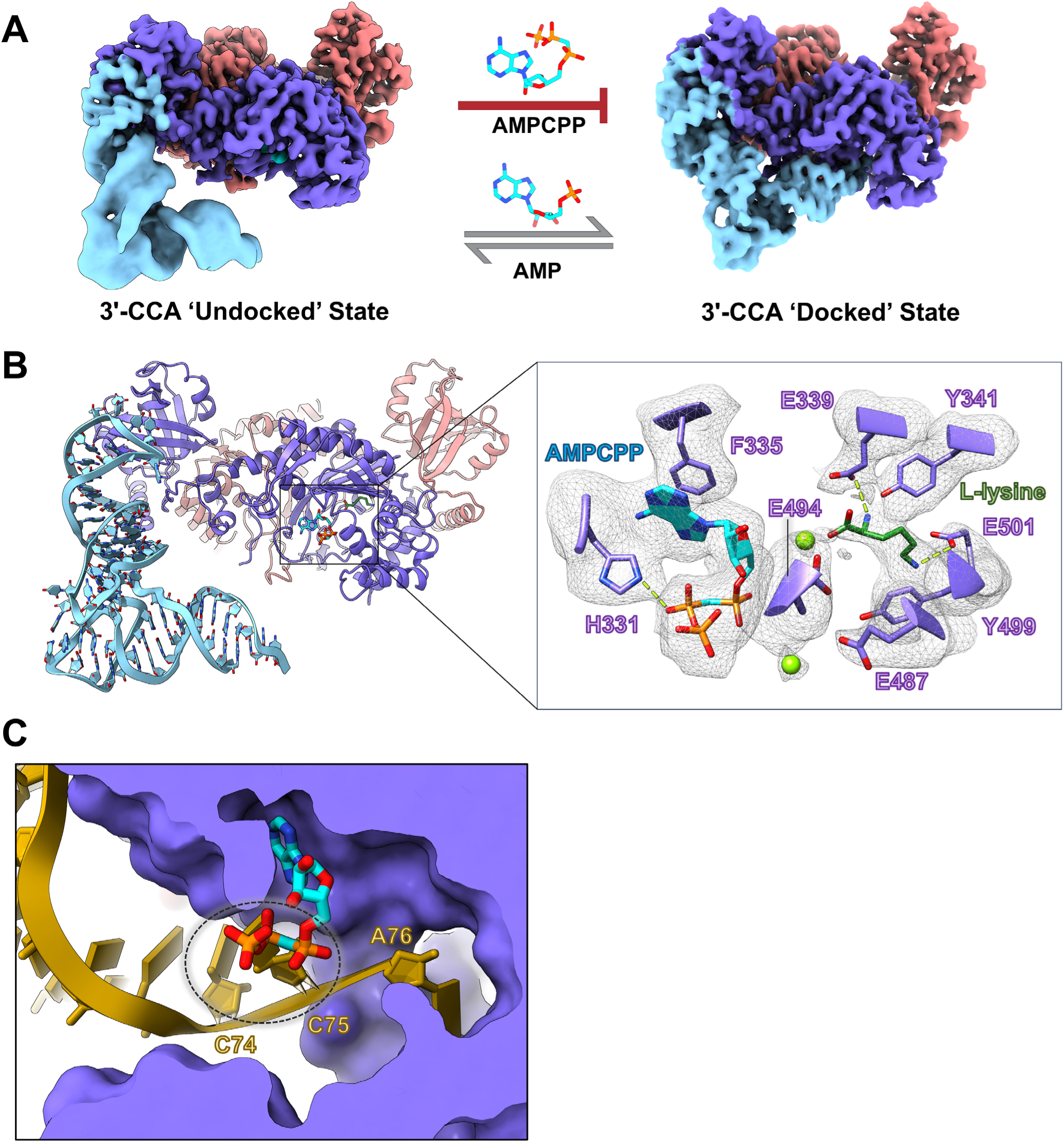
ATP state governs 3’-CCA end docking of tRNA^Lys3^ in the active site. (A) Cryo-EM reconstructions of LysRS bound to *in vitro* transcribed tRNA^Lys3^ (unmodified) in the presence of a non-hydrolysable ATP analog (AMPCPP) (left panel) and a post-hydrolysis ATP analog, AMP (right panel). The AMPCPP bound complex exists solely in the 3’-CCA ‘undocked’ conformation whereas the AMP bound complex can assume the ‘undocked’ and the ‘docked’ conformation. (B) Structural model for the AMPCPP bound ‘undocked’ state of the LysRS-tRNA^Lys3^ complex. The right inset shows the interactions of AMPCPP and L-lysine in the active site along with the corresponding cryo-EM map density. (C) The docked state of LysRS-tRNA^Lys3^ complex was aligned to the AMPCPP bound ‘undocked’ state. A cross-section view of the LysRS active site is shown highlighting the steric clash between the 3’-CCA end of tRNA^Lys3^ and the β- and γ-phosphate of ATP.

To understand why the non-hydrolyzable ATP analog, AMPCPP, prevented the complex from assuming the ‘docked’ state, we overlayed the structures for the docked and undocked states of the LysRS-tRNA^Lys3^ complex. This comparison revealed that the β- and γ-phosphates of the ATP analog clash with the 3’-CCA end of tRNA^Lys3^ in the LysRS active site (Figure 3C). Interestingly, H331 directly interacts with the β-phosphate of AMPCPP in the undocked state, and this same residue is involved in forming base-specific interactions with C75 of tRNA^Lys3^ in the docked state. Therefore, the docking of the 3’-CCA end of tRNA^Lys3^ cannot proceed until the lysyl-adenylate intermediate is formed and the cleaved pyrophosphate moiety is released from the active site. This is in contrast to the previously reported crystal structures for the docked state of yeast AspRS (y-AspRS)(40), a closely related class IIb AARS. The 3’-CCA end docking for tRNA^Asp^ in the y-AspRS active site was shown to be independent of the ATP state and it was postulated that the two steps in aminoacylation of tRNA^Asp^ are not ordered. Comparing our ‘docked’ state structure of the h-LysRS-tRNA^Lys3^ with that of y-AspRS-tRNA^Asp^ (PDB: 1ASZ) highlighted key differences in the active site architecture of these two enzymes. Compared to y-AspRS, the active site entrance in h-LysRS is more narrow (Supplementary Figure S4A) and is sterically blocked by the β- and γ-phosphate of ATP. h-LysRS encodes several insertions in the AAD (371–381, 388–396, 417–438) that are absent in y-AspRS, creating a narrow channel for insertion of the tRNA^Lys3^ 3’-CCA end (Supplementary Figure S4B-C). Thus, in contrast to AspRS, the two catalytic steps of aminoacylation in LysRS are structurally ordered. This ordering may have evolved to prevent conformational states that could act as catalytic dead-ends for the LysRS aminoacylation reaction.

### tRNA^Lys3^ modifications are directly recognized by h-LysRS and increase the affinity of the complex

tRNA^Lys3^ is heavily modified in the cell, especially in the loop regions, which alters its overall fold(37) compared to unmodified tRNA^Lys3^. Comparing the h-LysRS complexes with unmodified IVT tRNA^Lys3^ and modified cellular tRNA^Lys3^ allowed us to dissect the contributions of individual tRNA modifications to h-LysRS binding. A prime example of the crucial role of tRNA^Lys3^ modifications is the selective ordering of the N-terminal extension of h-LysRS in the presence of modified tRNA^Lys3^. In unmodified tRNA^Lys3^, U20 and U47 stack against the double-stranded helical core of tRNA^Lys3^, whereas in cellular tRNA^Lys3^ these bases are modified to D20 and D47, which are non-planar and assume an unstacked conformation (Figure 4A). The distance between the N3 atoms of U20 and U47 is ∼5 Å in the unmodified tRNA^Lys3^ whereas in modified tRNA^Lys3^, the distance between D20 and D47 is ∼22 Å. The wider groove created between the D-loop and variable loop by the unstacking of D20 and D47, along with the modified bases in the T-loop of modified tRNA^Lys3^ is crucial for the stable binding of the h-LysRS NTD (Figure 2A, 2B, and Supplementary Figure S4A).

**Figure 4.**
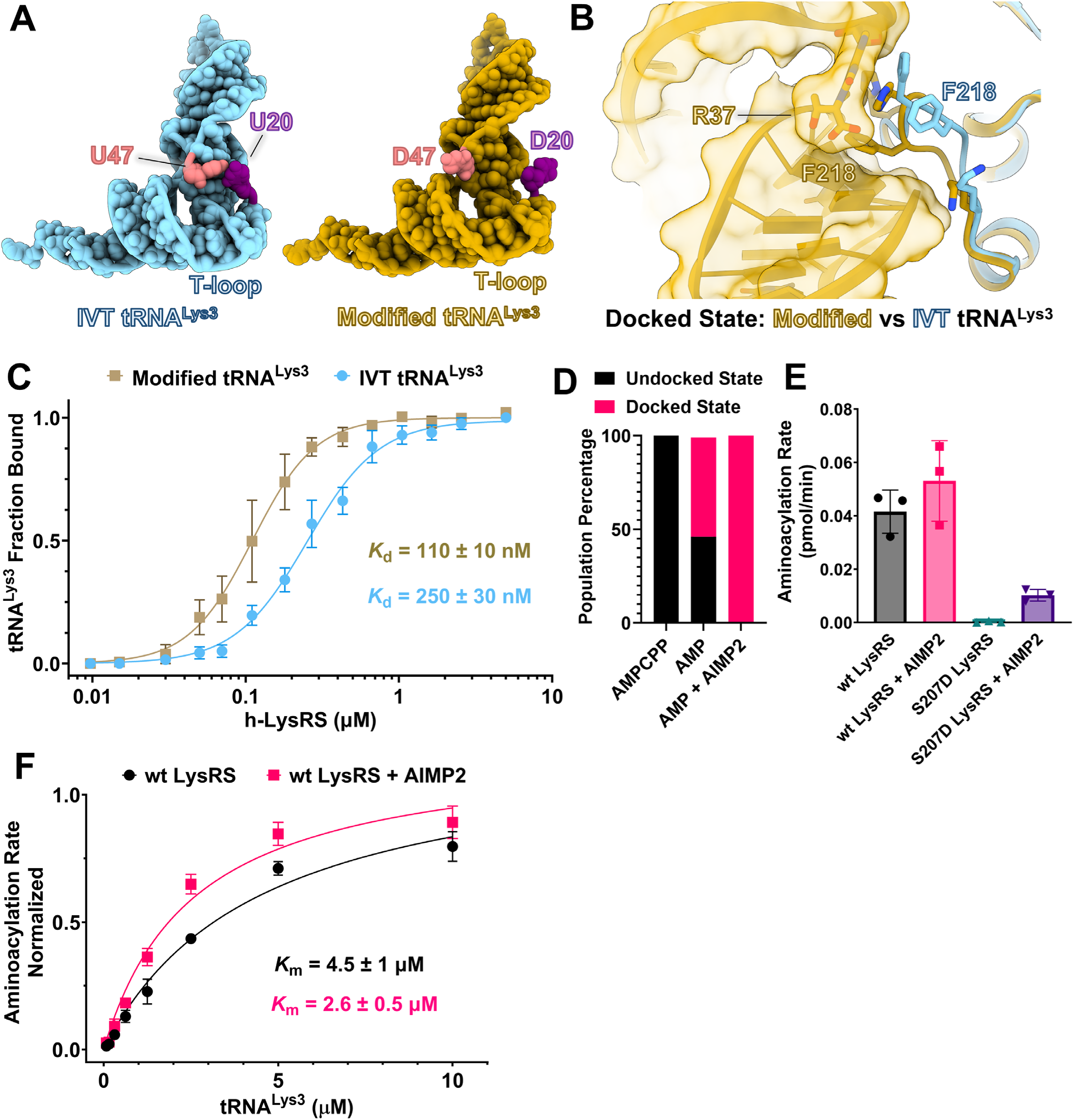
tRNA^Lys3^ modifications and AIMP2 enhance h-LysRS activity. (A) Cellular modified tRNA^Lys3^ (gold) and IVT unmodified tRNA^Lys3^ (sky blue) from their respective docked state structures are shown in cartoon representation with the bases shown as spheres. The nucleotide positions 20 (purple) and 47 (coral) in the modified and unmodified tRNA^Lys3^ are highlighted. (B) The docked state structures of LysRS with modified tRNA^Lys3^ (gold) and unmodified tRNA^Lys3^ (sky blue) were aligned and a magnified view of the linker region interacting with tRNA^Lys3^ is shown. Modified tRNA^Lys3^ (gold) is shown in surface view and unmodified tRNA^Lys3^ is hidden for clarity. LysRS (gold: modified tRNA^Lys3^ / sky blue: IVT tRNA^Lys3^) is shown in ribbon view. (C) EMSA titrations of h-LysRS binding to unmodified IVT tRNA^Lys3^ and modified tRNA^Lys3^ were carried out in triplicate. Fraction of tRNA^Lys3^ bound by h-LysRS was quantified and the mean is plotted against the protein concentration along with the associated standard error of the mean. The estimated binding affinity (*K*_d_) is denoted along with the associated error from the data fit. (D) The percentage of undocked and docked conformational states of the h-LysRS-tRNA^Lys3^ (IVT) complex are shown in bar chart form for the three cryo-EM datasets – h-LysRS-tRNA^Lys3^ in presence of AMPCPP and L-lysine, h-LysRS-tRNA^Lys3^ in presence of AMP and L-lysine, and h-LysRS-tRNA^Lys3^-AIMP2-N36 in presence of AMP and L-lysine. (E) The initial rates of wt and S2O7D h-LysRS aminoacylation in the presence and absence of AIMP2-N36 peptide, each measured in triplicate. The mean rates are plotted in bar chart form and error bars represent the associated standard deviation. (F) *K*_m_ measurement of wt h-LysRS for IVT tRNA^Lys3^ in presence and absence of AIMP2 peptide. The normalized aminoacylation rates are plotted against the tRNA^Lys3^ concentration. Each data point shown is the average of a triplicate measurement and the error bar corresponds to the associated standard deviation

The RNA modifications in the anticodon loop bases, S34 and R37, of cellular modified tRNA^Lys3^ directly interact with h-LysRS. R161 forms ionic interactions with the mcm modification of S34 whereas this interaction is missing for unmodified tRNA^Lys3^ (Supplementary Figure S4B). Similarly, the bulky N6 modification of R37 stacks against H217 and F218, and the ms^2^ moiety of R37 packs against H215 of the h-LysRS linker domain. H217 and H215 also form a hydrogen bonding network with the N6-threonyl group and the 5’-phosphate of R37, respectively. For unmodified tRNA^Lys3^, the h-LysRS linker region (208–220) adopts a significantly different conformation wherein H217 stacks against the A37 base but H215 and F218 do not participate in tRNA^Lys3^ binding (Figure 4B). Similarly, the hydrophobic interactions between m^2^G10 of cellular tRNA^Lys3^ with M212 and P214 of LysRS (Supplementary Figure S4C) result in a substantially more extensive protein-RNA interface and suggest that the affinity of h-LysRS for modified tRNA^Lys3^ would be higher than that for unmodified tRNA^Lys3^. Indeed, analyzing the population dynamics from our cryo-EM datasets showed that in the presence of AIMP2-N36 and modified tRNA^Lys3^, all the LysRS was present in the tRNA^Lys3^-bound form. In contrast, under identical experimental conditions in the presence of unmodified tRNA^Lys3^, almost half of the

LysRS was found in the apo-state (no tRNA^Lys3^ bound) (Supplementary Figure S4D and Supplementary Table S5). We carried out electrophoretic mobility gel-shift assays (EMSA) of h- LysRS with cellular modified tRNA^Lys3^ and IVT unmodified tRNA^Lys3^ to probe the effect of tRNA^Lys3^ modifications on binding affinity. Indeed, EMSA binding assays confirmed that the affinity of h-LysRS for fully modified tRNA^Lys3^ was ∼2.5-fold higher than for IVT tRNA^Lys3^ (Figure 4C and Supplementary Figure S4E).

### AIMP2 drives the equilibrium of the h-LysRS-tRNA^Lys3^ complex towards the ‘docked’ state

To understand the effect of AIMP2 binding on the conformational landscape of the LysRS- tRNA^Lys3^ complex, we collected another cryo-EM dataset for the LysRS-tRNA^Lys3^ (IVT) complex in the presence of AMP, L-lysine, and AIMP2-N36 peptide. This dataset yielded two reconstructions – apo-LysRS and LysRS-tRNA^Lys3^ complex. Strikingly, in the presence of AIMP2- N36, all the LysRS-tRNA^Lys3^ complex existed in the ‘docked’ state. For an identical experimental setup, except for the presence of AIMP2, the LysRS-tRNA^Lys3^ complex exhibited an equilibrium between the ‘docked’ (55%) and the ‘undocked’ (45%) states (Figure 4D, Supplementary Table S5). This indicates that AIMP2 binding allosterically drives the LysRS-tRNA^Lys3^ complex towards the ‘docked’ state, suggesting an active role in enhancing the catalytic efficiency of LysRS.

To test our structure-based predictions that AIMP2 may enhance the aminoacylation activity of LysRS, we carried out aminoacylation assays of LysRS in the absence and presence of AIMP2-N36. Interestingly, AIMP2-N36 was able to rescue the aminoacylation activity in the catalytically inactive S207D mutant of LysRS (Figure 4E). The presence of AIMP2-N36 also showed a modest increase in the aminoacylation rate of wt LysRS and despite the rescue effect on the catalytic activity of S207D LysRS, wt LysRS exhibited ∼5-fold higher activity than S207D LysRS. We also measured the *K*_m_ of wt LysRS for tRNA^Lys3^ in the absence and presence of AIMP2-N36 and found that in the presence of AIMP2, the *K*_m_ of wt LysRS for tRNA^Lys3^ decreased by two-fold (Figure 4F). These results corroborate our observations from cryo-EM studies and indicate that AIMP2 and the MSC play an active role in tRNA^Lys3^ aminoacylation by facilitating the transition of the h-LysRS-tRNA^Lys3^ complex towards the ‘docked’ state and enhancing the *K*_m_ of h-LysRS for tRNA^Lys3^.

## DISCUSSION

tRNA aminoacylation is a fundamental biological process across all living organisms and AARS are proposed to have been among the first enzymes in the evolutionary timeline(41). Despite exhibiting a high degree of conservation, the aminoacylation machinery in higher eukaryotes has acquired several additional layers of regulation and complexity over the course of evolution. For example, most prokaryotes carry 20 AARS, one for every aa, but in humans, two distinct sets of AARS are employed for carrying out cytosolic and mitochondrial aminoacylation reactions. Furthermore, almost half of the cytosolic AARS are housed in the MSC where they presumably carry out their canonical aminoacylation function. In humans, AARS also carry out secondary functions in physiological processes ranging from regulation of nuclear transcription to immune responses(42). Although the simpler bacterial AARS enzymes have been well-characterized, a structural and mechanistic understanding of the more complex aminoacylation machinery in humans and other higher eukaryotes has been lacking.

h-LysRS is an indispensable component of the cellular translation machinery, is a regulator of nuclear transcription(15,16), and is also a critical host factor for the HIV-1 life cycle(32). Despite its myriad roles, the structural and mechanistic basis for recognition and aminoacylation of tRNA^Lys^ has been lacking. Furthermore, the roles of the MSC and tRNA modifications in the function of AARS represent fundamental gaps in our understanding of aminoacylation in humans. Our structure of h-LysRS bound to cellular modified tRNA^Lys3^ and a peptide derived from the MSC scaffold protein AIMP2 is the first structure of a human AARS bound to a fully modified cellular tRNA and its MSC interacting partner (Figure 1B-C).

The tRNA^Lys3^ modifications play an integral role in mediating the specificity of h-LysRS for tRNA^Lys3^, and directly modulate the structure of the h-LysRS-tRNA^Lys3^ complex (Figure 1, Figure 2, and Supplementary Figure S4A-C). The eukaryotic specific NTD of h-LysRS (1–65) folds into an extended helix in the presence of modified tRNA^Lys3^ and forms an extensive interface with the tRNA^Lys3^ acceptor arm, D-loop, variable loop, and T-loop (Figure 2A). The tRNA^Lys3^ modifications are essential for ordering the h-LysRS NTD, which remains disordered in the presence of unmodified IVT tRNA^Lys3^ (Figure 3A-B). Previous characterization of rabbit tRNA^Lys^ species identified three major isoforms, designated tRNA^Lys1^, tRNA^Lys2^ and tRNA^Lys3^ based on the order in which they eluted from an anion exchange column(19). The sequence and RNA modifications observed in rabbit tRNA^Lys3^ are conserved in human tRNA^Lys3^ investigated here. Comparing the RNA modifications between the three rabbit tRNA^Lys^ isoforms shows that almost all the modifications in the D-loop, variable loop, and T-loop of tRNA^Lys3^ that mediate interaction with h-LysRS are conserved in tRNA^Lys1^ and tRNA^Lys2^. A notable difference between the three isoforms is the lack of S34 modification in tRNA^Lys1^ and tRNA^Lys2^.

Human AspRS and AsnRS, the two other members of the class IIb AARS family, have also evolved an NTD that is absent in their bacterial counterparts. However, whether the NTDs of h-AspRS and h-AsnRS function similarly to h-LysRS remains unresolved due to the lack of sequence conservation in this region (Supplementary Figure S5A). Alphafold predictions for the secondary structure profile of the NTD of h-LysRS, h-AspRS, and h-AsnRS vary significantly, whereas their ABD domains fold into a highly conserved architecture (Supplementary Figure S5B). Furthermore, the effect of the unique sequence and RNA modification profiles of tRNA^Asp^ and tRNA^Asn^ on their interactions with h-AspRS and h-AsnRS remains uncharacterized. Further structural studies on h-AspRS and h-AsnRS are needed to understand the roles of their respective NTDs and identify commonalities across the class IIb family of cytosolic AARS. Recently, the human mitochondrial tRNA maturation pathway was structurally characterized(43), and it revealed that the mitochondrial tRNA methyltransferase 10C (TRMT10C) employs an extended α-helical motif that is remarkably similar to h-LysRS NTD in its charge distribution and interaction network with tRNA. This suggests that the h-LysRS NTD may be a conserved interaction motif in metazoan tRNA binding proteins.

The evolutionary advantage of housing cytosolic AARS in the MSC has been unclear and several potential functions have been proposed for the MSC(11,44). h-LysRS association with the MSC has been proposed to protect the aminoacylation function by preventing modification and/or recruitment into alternate cellular pathways(45). Herein, we show that the association of h-LysRS with the MSC via the scaffold protein AIMP2 enhances the aminoacylation efficiency of tRNA^Lys3^ (Figure 1 and 4). AIMP2 allosterically modulates the equilibrium of the h-LysRS- tRNA^Lys3^ complex towards the ‘docked’ state (Figure 5). Whether an association with the MSC enhances the catalytic activity of other AARS remains to be explored.

**Figure 5.**
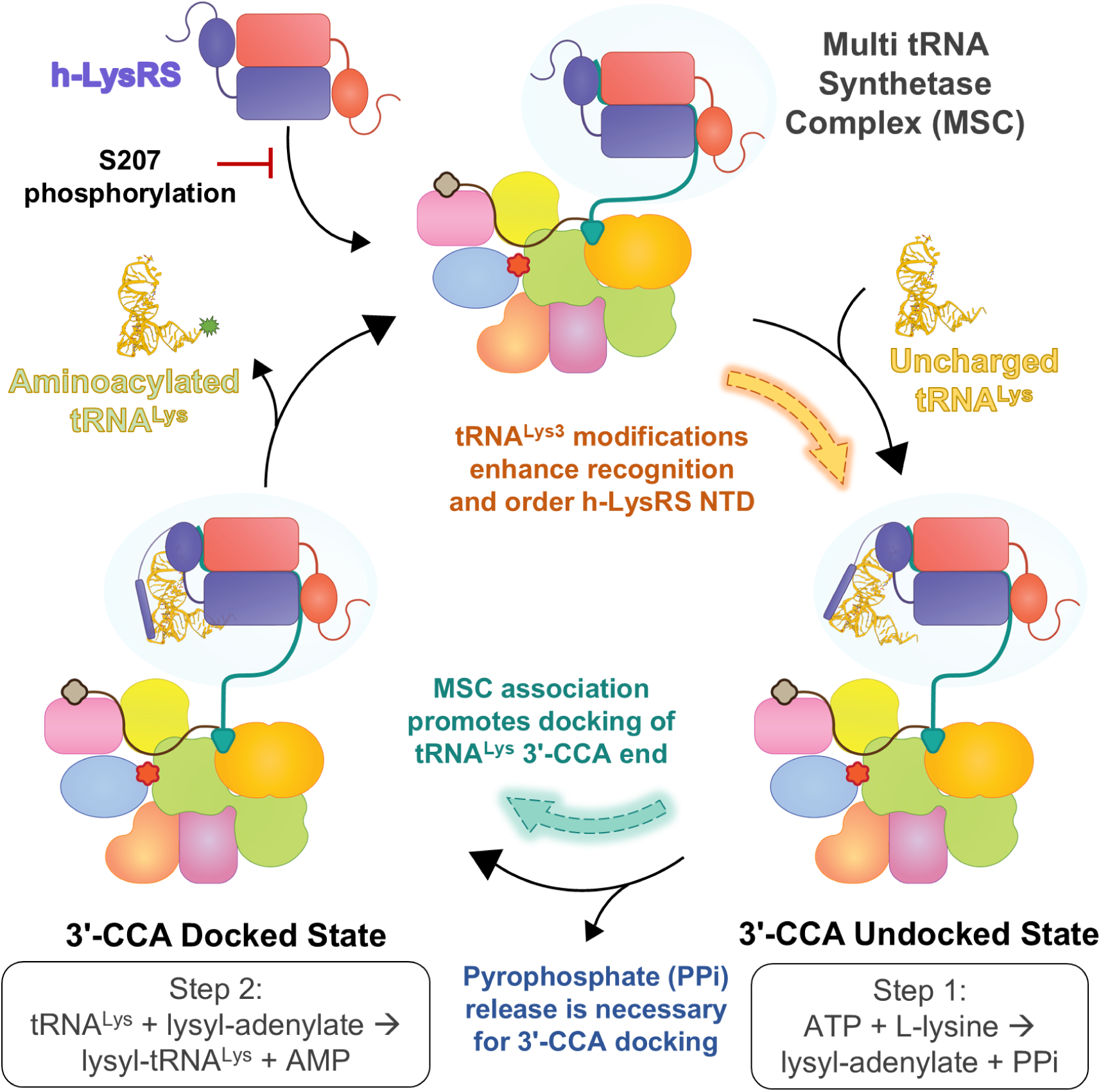
Schematic model for tRNA^Lys3^ aminoacylation and the roles of tRNA^Lys3^ modifications and the MSC in h-LysRS function. h-LysRS associates with the MSC via the scaffold protein AIMP2. Phosphorylation at S2O7 residue of h-LysRS prevents association with the MSC. h-LysRS selectively binds tRNA^Lys^ and the fidelity of this interaction is imparted by the anticodon loop and the unique tRNA^Lys^ modifications. tRNA^Lys^ modifications are also critical in ordering the h-LysRS NTD into an extended helix that docks at the base of the T-loop and interacts along the length of the anticodon arm. The two catalytic steps in tRNA^Lys^ aminoacylation are structurally ordered by h-LysRS active site, β and γ-phosphate of the bound ATP create a steric block and prevent the docking of tRNA^Lys^ 3’-CCA end. In the ATP-bound state, the h-LysRS-tRNA^Lys^ complex exists exclusively in the ‘3’-CCA undocked’ state. Once the ATP hydrolysis catalyzed lysyl-adenylate intermediate is formed and the pyrophosphate (PP_i_) byproduct is released, tRNA^Lys^ 3’-CCA end can be docked in the active site. In the 3’-CCA docked state, the 3’-A76 nucleotide is unstacked and flipped into the active site, positioning the 3’-hydroxyl of the A76 base for attack on the lysyl-adenylate intermediate. Association with the MSC allosterically drives the equilibrium of the h-LysRS-tRNA^Lys^ complex from the ‘undocked’ state towards the ‘docked’ state, thereby enhancing the catalytic efficiency of h-LysRS. After the completion of the second step in tRNA^Lys^ aminoacylation, aminoacylated tRNA^Lys^ is released along with the byproduct AMP, and the catalytic cycle can begin again.

Aminoacylation by all AARS is a two-step catalytic reaction, wherein the cognate aa is first activated using ATP hydrolysis to generate the aminoacyl-adenylate intermediate and then the activated aa is conjugated to the 3’-end of the respective tRNA (2’-hydroxyl for class I AARS and 3’-hydroxyl for class II AARS). These two catalytic steps in tRNA^Lys3^ aminoacylation are sequentially ordered by the LysRS active site such that the docking of the tRNA^Lys3^ 3’-CCA end in the active site can only occur after ATP hydrolysis driven activation of L-lysine is completed (Figure 3). This is in contrast to the closely related AspRS, wherein the active site can simultaneously bind ATP, L-aspartic acid, and the 3’-CCA end of tRNA^Asp^ (Supplementary Figure S3). This stems from the differences in the active site architecture wherein insertions between motifs 2 and 3 of h-LysRS lead to an active site entrance that is much narrower than AspRS and incompatible with simultaneous binding of ATP and the 3’-CCA end of tRNA^Lys3^. AsnRS is believed to have evolved from AspRS(46), and its active site is more similar to AspRS than LysRS, suggesting that the two catalytic steps in tRNA^Asn^ aminoacylation are likely not ordered.

Recently, several neurological and metabolic disorders have been linked to dysregulation of tRNA modifications, collectively referred to as tRNA modopathies(47). For example, several tRNA^Lys3^ modopathies including microcephaly(48), nephropathy(49), diabetes(50), and cancer(51) are linked to downregulation of the S34 and R37 modifications. The exact mechanistic basis for how changes in the tRNA modification landscape lead to these disorders remains unclear and understudied. Our results herein show the critical role of tRNA^Lys3^ modifications in mediating the specificity of h-LysRS-tRNA^Lys3^ interaction and predict that downregulation of these modifications, although not essential for *in vitro* tRNA binding and aminoacylation, would likely impair the aminoacylation efficiency of tRNA^Lys^ in cells, which may impact the translation of lysine-rich proteins. Modifications may also provide a fitness advantage under certain environmental conditions including cellular stress(52,53). The significant structural rearrangements observed with modified tRNA^Lys3^ bound to LysRS may finetune the translation machinery under specific cellular conditions. For example, during T cell activation, ms^2^t^6^A37 in the anticodon loop of tRNA^Lys3^ showed dynamic changes with an impact on frameshifting(54).

In summary, our results provide an in-depth understanding of tRNA^Lys3^ recognition and aminoacylation by h-LysRS and delineate the critical roles of tRNA^Lys3^ modifications and the MSC in h-LysRS function. The structural and mechanistic insights presented herein provide a strong foundation for future work aimed at understanding the aminoacylation machinery in humans as well as the increasing number of neurological and metabolic disorders linked to defects in the human aminoacylation and tRNA modification machinery(4,25–27).

## MATERIALS AND METHODS

### Expression and purification of wt and S207D LysRS

Full-length LysRS (wt and S207D) was cloned into pET11a protein expression vector with an N-terminal 6X-His affinity tag and recombinantly expressed in *Escherichia coli* BL21 (DE3) cell line. Cells were grown in Terrific Broth to an O.D. of 0.7-0.8 and then induced with 0.4 mM isopropyl β-D-1-thiogalactopyranoside (IPTG) at either 18°C for 16 h. Cells were harvested via centrifugation, resuspended in lysis buffer [50 mM Tris-Cl pH 7.5, 500 mM NaCl, 5% glycerol (v/v), 0.5 mM TCEP (tris(2-carboxyethyl)phosphine), protease inhibitor cocktail tablets, at 4°C], and subsequently lysed using a microfluidizer. The lysate was clarified by centrifugation to remove bacterial cell debris, and the supernatant was subjected to PEI (polyethyleneimine) precipitation at 0.45% w/v to remove contaminating nucleic acids. The supernatant from the PEI precipitation step was then used for ammonium sulfate precipitation (55% saturation) to separate the protein fraction from the contaminants. Pellets from ammonium sulfate precipitation were then solubilized in Buffer A [50 mM Tris-Cl pH 7.5, 300 mM NaCl, 5% glycerol (v/v), 0.5 mM TCEP, at 4°C ] and applied to a Ni-NTA resin packed in a gravity-flow column. The column was washed extensively with Buffer A containing 20 mM imidazole and then 6X-His tagged LysRS was eluted using Buffer A containing 400 mM imidazole. Eluted protein was dialyzed overnight into Buffer B [10 mM potassium phosphate pH 7.5, 150 mM KCl, 5% glycerol (v/v), 0.2 mM TCEP, at 4°C ]. Dialyzed eluate from Ni-NTA column was then applied to a hydroxyapatite column (BioRad) and subjected to a gradient of increasing phosphate concentration [Buffer C - 400 mM potassium phosphate pH 7.5, 150 mM KCl, 5% glycerol (v/v), 0.2 mM TCEP, at 4°C] to further enrich the LysRS fraction. Peaks from the hydroxyapatite column run were run on an SDS-PAGE gel and fractions containing LysRS with purity >95% were then pooled and dialyzed into protein storage Buffer C [50 mM HEPES pH 7.5, 150 mM NaCl, 5% glycerol (v/v), 0.2 mM TCEP]. Purified LysRS was then concentrated using Amicon® Ultra-15 Centrifugal Filter Concentrators with a 10 kDa cutoff, aliquoted, flash frozen in liquid nitrogen, and stored at -80°C .

### Purification of cellular modified tRNA^Lys3^

The protocol to purify fully-modified human tRNA^Lys3^ using an affinity-capture-based method was adapted from Drino et al.(55) HEK293T (ATCC) or Jurkat cell lines (NIH AIDS Reagent Program) were cultured at 5% CO_2_ in 10% FBS (v/v) and 5% penicillin/streptomycin (Gibco) supplemented DMEM or RPMI (Gibco), respectively. Cells were washed with phosphate-buffered saline (PBS) (Thermo Fisher Scientific) and lysed with TRIzol (Invitrogen) per the manufacturer’s instructions. The total cellular RNA was extracted with chloroform by centrifuging at 16,000x *g* for 15 min at 4°C, and the aqueous layer was transferred to a fresh tube. LiCl stock solution (7.5 M) was added to the total RNA solution to a final concentration of 4 M, and the mixture was incubated overnight at –20°C, followed by centrifugation at 13,700x *g* for 10 min at 4°C. This step allows for precipitation of high molecular weight RNA. The supernatant enriched for small RNAs was collected and ethanol precipitated with 2 µL of 5 mg/ml glycogen. The pellet was recovered and dissolved in 500 µL of ion exchange wash buffer (20 mM Tris, pH 7.5, 10 mM KCl, 1.5 mM MgCl_2_, 50 mM NaCl). The sample was loaded onto a HiTrap Q FF anion exchange column (5 ml) (Cytiva) on an ÄKTA pure chromatography system (Cytiva) equilibrated with the same buffer. RNA was eluted using a NaCl gradient from 10 – 1000 mM at a flow rate of 0.3-0.5 ml/min. Fractions eluting between 400-600 mM NaCl were collected and immediately precipitated with isopropanol at -20°C and dissolved in milliQ-H_2_O.

A human tRNA^Lys3^-specific affinity column was generated by coupling a 5′ amino-modified DNA oligonucleotide complementary to the 5′ 36 nt of tRNA^Lys,3^ (5′ AAAAGTCTGATGCTCTACCGACTGAGCTATCCGGGC 3′) purchased from IDT, to a HiTrap NHS-activated HP column (1 ml) (Cytiva) through a stable amide linkage as described.(55) Briefly, the affinity oligonucleotide stock solution was ethanol precipitated and dissolved in 1 ml of coupling buffer (0.2 M NaHCO_3_, 0.5 M NaCl, pH 8.3). The NHS-activated HP column was washed with 2 ml of 1 mM ice-cold HCl three times at a maximum flow rate of 0.5 ml/min. The oligonucleotide solution was immediately injected onto the column and the column was sealed and incubated for 30 min at 25°C (or 4 h at 4°C). Uncoupled activated groups were deactivated and unbound oligonucleotides were removed by repeated washings with washing buffer A (0.5 M ethanolamine, 0.5 M NaCl, pH 8.3) and washing buffer B (0.1 M NaOAc, 0.5 M NaCl, pH 4). The coupling efficiency of the oligonucleotide to the column was ∼40% and the column was stored in 0.05 M Na_2_HPO_4_, 0.1% NaN_3_, pH 7 at 4°C.

To capture human tRNA^Lys,3^, the small RNA pool was denatured by incubation at 75°C for 3 min, followed by snap-cooling on ice. The RNA was mixed with 20 ml of precooled binding buffer (30 mM HEPES-KOH, pH 7.0, 1.2 M NaCl, 10 mM MgCl_2_). The affinity column was equilibrated with the binding buffer and binding of tRNA to the column was performed under controlled temperature conditions – the RNA pool was circulated through the affinity column for 90 min at 65°C, and then the temperature was gradually decreased to 40°C over a 3 to 4 h period. The column was washed with 10 ml of 2.5 mM HEPES-KOH, pH 7.0, 0.1 M NaCl, 10 mM MgCl_2_ at 40°C. Finally, tRNA^Lys3^ was eluted from the column with 0.5 mM HEPES-KOH, pH 7.0 and 1 mM EDTA at 75°C. The eluted RNA was immediately ethanol precipitated and the purified tRNA^Lys3^ was checked on a 10% denaturing polyacrylamide gel. The yield of modified human tRNA^Lys3^ starting with around 100 million HEK 293T cells was ∼5 µg.

### LC-MS validation of tRNA^Lys3^ modifications

To detect tRNA^Lys3^ modifications, liquid chromatography tandem mass spectrometry (LC-MS/MS) was performed. The RNA sample was digested using 50 U/µg of commercially purchased RNase T1 (Worthington Biochemical, Lakewood, NJ) in 110 mM ammonium acetate at 37 °C for 1.5 h. The enzymatic digestion products were separated using a Dionex Ultimate 3000 (Thermo Scientific) UHPLC system equipped with an Xbridge BEH C18 column (2.1 mm x 150 mm, 2.5 μm, Waters) using ion-pair reverse phase liquid chromatography (IP-RPLC). Mobile phases were 8 mM triethylamine (TEA) and 200 mM hexafluoroisoproponal (HFIP) in water (mobile phase A: MPA) and 8 mM TEA and 200 mM HFIP in 50:50 water: methanol (mobile phase B: MPB). The IP-RPLC gradient profile was 5% MPB until 5 min followed by a linear ramp to 73% MPB at 40 min. The column was flushed with 100% MPB for 5 min and re-equilibrated for 15 min at 5% MPB. The flow rate was 80 µL/min with a column temperature of 60°C.

An LTQ-XL linear ion trap mass spectrometer (Thermo Fisher Scientific) was used to detect the resolved digestion products in negative polarity mode. Electrospray ionization conditions were as follows: capillary temperature 375°C, capillary voltage of -30 V, spray voltage of 3.7 kV and 35, 20, and 20 arbitrary flow units of sheath, auxiliary and sweep gas, respectively. Data was acquired with the range of *m/z* 600-2000; MS/MS scans were collected through data-dependent acquisition of the four most abundant MS ions. The maximum injection was fixed at 250 ms with an AGC cutoff of 500 counts. LC-MS/MS data analysis was performed using RNAModMapper (RAMM)(56,57) and Xcalibur v4.1. (Thermo Fisher Scientific). All expected RNase T1 digestion products from human tRNA^Lys3^ were detected in the mass spectral data and could be mapped to the tRNA sequence (Supplementary Figure S1A and Supplementary Table S1). These digestion products included the known modifications on human tRNA^Lys3^, *N*^2^- methylguanosine at position 10 (m^2^G10), dihydrouridine at positions 16 and 20 (D16, D20), 5- methoxycarbonylmethyl-2-thiouridine at position 34 (mcm^5^s^2^U34), 2-methylthio-*N*^6^- threonylcarbamoyladenosine at position 37 (ms^2^t^6^A37), *N*^7^-methylguanosince at position 46 (m^7^G46), 5-methylcytidine at position 49 (m^5^C49), 5,2’-*O*-dimethyluridine at position 54 (T^m^54), and 1-methyladenosine at position 58 (m^1^A58). Automated analysis of MS/MS fragmentation data by RAMM(57) confirmed the sequences of all digestion products except the fully modified anticodon sequence ‘ACU[mcm^5^s^2^U]UU[ms^2^t^6^A]AYCUGp’ and the variable loop digestion product ‘[m^7^G]U[m^5^C][m^5^C]AGp’. These two fragments were analyzed and verified manually. One limitation of the mass spectrometry-based approach is that uridine and pseudouridine cannot be differentiated. Thus, the presence of pseudouridine at positions 27, 39 and 55 (ψ27, ψ39, ψ55) was not confirmed by LC-MS. Overall, the LC-MS analysis identified nine RNA modifications including anticodon loop modifications known to be present in human tRNA^Lys3^ in the purified tRNA sample.

### In vitro transcription of tRNA^Lys3^

Human tRNA^Lys3^ was *in vitro* transcribed by in-house purified T7 RNA polymerase(58) prepared using plasmid p6XHis-T7(P266L),(59) a gift from Anna Pyle (Addgene plasmid #174866). The plasmid template used for transcription (pIDTSmart-htRNALys3)(60) was digested with FokI followed by buffered phenol-chloroform extraction and ethanol precipitation.(61) Transcription reactions (4 mL) containing 240 µg linearized plasmid template or PCR product from a 1-mL PCR reaction were performed in 80 mM HEPES, pH 8, 30 mM Mg(OAc)_2_, 10 mM dithiothreitol (DTT), 5 mM spermidine, 0.01% Triton-X-100, 4% PEG8000, 0.5 mg/mL BSA, 4 mM of each rNTP, 10 µg/mL pyrophosphatase and an optimized amount of recombinant P266L T7 RNA polymerase at 37°C for 5 h. The reaction was quenched with 50 mM EDTA, pH 8, and the tRNA was purified by acid-phenol chloroform extraction followed by anion exchange chromatography as described.(62,63) Briefly, the tRNA product from a 4-mL transcription was loaded onto a 3-mL DEAE Sepharose Fast Flow gravity column (Cytiva) equilibrated with 20 mM Tris-HCl, pH 8.0, 150 mM NaCl, and 5 mM MgCl_2_ and washed with the same buffer containing 250 mM NaCl. Bound tRNA molecules were eluted from the column using 15 mL of 1 M NaCl buffer. Fractions containing the tRNA were identified by urea-PAGE, pooled, and ethanol precipitated. tRNAs were resuspended in MilliQ-H_2_O and the purity was assessed by urea-PAGE.

### Cryo-EM grid preparation and data collection

Purified wt LysRS was mixed with either cellular purified tRNA^Lys3^ or IVT tRNA^Lys3^ in a 1:1 stoichiometric ratio and a final concentration of 10 µM h-LysRS-tRNA^Lys3^ along with 2.5 mM L-lysine and 2 mM of either AMP (adenosine monophosphate) or AMPCPP (α,β- methyleneadenosine-5’-triphosphate). For datasets containing AIMP2-N36, the chemically synthesized peptide was added to a final concentration of 100 µM. Complex formation was allowed to proceed for 60 min at 4°C. The assembled complex (4 µL) was applied to C-flat R2/1 holey carbon copper grids (EMS) after glow discharging the grids at 11 mA for 30 sec (Pelco). Grids were blotted at blot force 1 for 6 sec and plunge frozen in liquid ethane using the Vitrobot Mark IV system (Thermo Fisher Scientific) maintained at 10°C. The detergent CHAPSO (0.075% final concentration) was added just before grid freezing to ensure uniform ice thickness and particle distribution.

Grids were screened for optimal ice thickness and particle concentration. Data collection was carried out from a single screened grid using a 300 kV Titan Krios cryo-transmission electron microscope (Thermo Fisher Scientific) equipped with a K3 camera (Gatan) and an imaging energy filter (Gatan) operated at a slit width of 15 eV. The dataset was collected in counting super-resolution mode with a nominal magnification of 81,000x leading to a physical pixel size of 1.07 Å (super-resolution pixel size is 0.535 Å). The data were collected at a dose rate of 15 e^-^/pixel/sec with a total electron dose of 50 e^-^/Å^2^ applied over 40 frames and a defocus range of - 1.0 µm to -2.0 µm.

### Cryo-EM data processing and model building

The cryo-EM data processing workflow was carried out using cryoSPARC v4(64). Movie frames were motion corrected using Patch motion correction and binned two-fold to yield a stack of motion-corrected micrographs with a pixel size of 1.07 Å. These micrographs were then manually curated, and an initial round of particle picking was performed using Blob picker on a subset of 500 micrographs. A 2D classification job on these initial particle stack yielded ideal 2D templates that were used to perform template-based particle picking on the entire dataset. The particle stack from template picker was further curated to remove damaged/denatured particles using 2D classification. After 2D classification, selected particles were carried forward to *Ab initio* reconstruction for generating three different volumes and used as 3D references for heterogeneous refinement of the particle stack. Particles corresponding to apo-LysRS, LysRS- tRNA^Lys3^ undocked state, and LysRS-tRNA^Lys3^ docked state were segregated and further refined using Non-Uniform (NU) refinement. These particle stacks were further curated using 3D classification to improve compositional and conformational homogeneity. The final particle stacks were then polished using Reference-based motion correction and used for NU-refinement to create the final reconstructions. All datasets were processed as above, and the details are included in Tables S2-S5. The final reconstructions were uploaded to the Electron Microscopy Data Bank (EMDB).

Model building was done in Coot(65) with a previously published model of human LysRS bound to AIMP2 peptide (PDB: 6ILD) and the crystal structure of mammalian tRNA^Lys3^ (PDB: 1FIR) as the starting templates. After initial rigid-body docking, the tRNA^Lys3^ and LysRS residues were manually fit into the high-resolution map using real-space refinement modules. Some regions of LysRS that were missing in the starting template were built *de novo* into the high-resolution map. Two rounds of real-space refinement in PHENIX(66) were carried out and the outliers were manually corrected in Coot before deposition in the Protein Data Bank (PDB).

### Aminoacylation Assays

*In vitro* transcribed or fully modified cellular tRNA^Lys3^ was folded in 50 mM HEPES, pH 7.5 by heating at 80°C for 2 min, 60°C for 2 min, followed by the addition of 10 mM MgCl_2_, incubation for 5 min at room temperature and placement on ice. The active tRNA concentration was determined by aminoacylation assays as described below, with 1 µM LysRS and time points taken up to 40 min to ensure plateau-level charging.

AIMP2 N36 peptide(67) purchased from Biosynth was resuspended in 1 mM TCEP, 10% glycerol and stored at -80°C. Concentration was determined by measuring the absorbance at 280 nm using an extinction coefficient of 5240 M^-1^ cm^-1^. Aminoacylation assays for determination of kinetic parameters in the presence and absence of AIMP2 N36 peptide were performed as follows. LysRS was diluted to 200 nM in 50 mM HEPES, 0.1 mg/mL BSA, 20 mM β- mercapt<u>oethanol</u> (βME), and 20 mM KCl with or without 50-fold molar excess AIMP2 N36 peptide and incubated on ice for ∼1 h prior to starting the reaction. Aminoacylation was conducted at 30°C in the presence of 0.08-10 µM active unmodified tRNA, refolded as described above, 0.1 mg/mL BSA, 20 mM KCl, 20 mM MgCl_2_, 20 mM βME, 4 mM ATP, 50 mM HEPES, pH 7.4, 20 µM lysine, and 2.5 µM [^3^H]lysine (60 Ci/mmol in 2% ethanol, American Radiolabeled Chemicals). Reactions were initiated with 10 nM LysRS. At 20 sec time intervals out to 2 min, an aliquot (10 µL) of reaction mix was quenched on Whatman 3mM filter pads presoaked with 5% trichloroacetic acid (TCA). Pads were washed with excess 5% ice-cold TCA three times for 20 min each, before drying and analysis by liquid scintillation counting. Specific activity was calculated with 1x reaction buffer (no washes) to calculate [^3^H] counts per min per pmol lysine.

### Electrophoretic mobility shift assays (EMSA)

EMSAs were conducted by incubating increasing concentrations of wt LysRS (0 to 5 µM) with 37.5 nM unmodified or modified tRNA^Lys3^ in binding buffer (20 mM HEPES pH 7.4, 60 mM NaCl, 10 mM KCl, 1 mM MgCl_2_, 400 µM DTT, 8% glycerol) at room temperature for 30 min. Complexes were run on 4-16% NativePAGE gels (Invitrogen) in a Tris-Borate native buffer (pH 8.3) for 2 h at 100V, stained with SYBRgold (Invitrogen) and imaged using Cy3 filters on an Amersham™ Typhoon™ RGB imager (Cytiva). Binding was quantified using ImageJ software and the proportion of bound to total tRNA^Lys3^ was calculated. Dissociation constants (*K*_d_) were calculated by plotting percent bound tRNA^Lys3^ versus LysRS concentration and fitting the data to the Hill equation(68) using GraphPad Prism software version 10.3.0 for MacOS.

## ACKNOWLEDGEMENTS AND FUNDING

This work was funded by the National Institutes of Health (NIH) grants R21 AI157890 (to Y.X.), NIH GM058853 (to P.A.L.), T32 GM141955 (to C.R.B.), R35 GM141880 and R01 AI150493 (to K.M.F.).

## AUTHOR CONTRIBUTIONS

S.C.D., K.M.F., Y.X. conceptualized and designed the study. S.C.D. carried out the cryo-EM sample preparation, data collection, data processing, and model building. C.P. and A.K. purified the cellular modified tRNA^Lys3^. S.C.D. and C.R.B. purified the wt and S207D LysRS.

C.R.B. carried out the aminoacylation assays and EMSAs. C.H. and P.A.L. carried out the LC-MS validation and analysis. S.C.D., Y.X., and K.M.F. wrote the original draft of the manuscript. All authors contributed to review and editing.

## RESOURCE AVAILABILITY

Plasmids and cell lines used in this study are available upon request. Any further information and requests for resources should be directed to and will be fulfilled by S.C.D.

## DATA AVAILABILITY

The coordinates and cryo-EM density maps for all the structures reported here have been deposited in the PDB and EMDB, respectively. The accession codes are as follows - h-LysRS-AIMP2-modified tRNA^Lys3^ (Docked State, AMP bound) complex (PDB: 9DPL, EMD-47106); h-LysRS-IVT tRNA^Lys3^ (Undocked State, AMPCPP bound) complex (PDB: 9DPB, EMD-47101); h-LysRS-IVT tRNA^Lys3^ (Undocked State, AMP bound) complex (PDB: 9DOW / EMD-47094); h-LysRS-IVT tRNA^Lys3^ (Docked State, AMP bound) complex (PDB: 9DPA, EMD-47100).

## SUPPLEMENTARY DATA STATEMENT

Supplementary Data is available online.

