## Supplementary Data for "Structural basis for aminoacylation of cellular modified tRNA^Lys3^ by human lysyl-tRNA synthetase"

### Figure S1

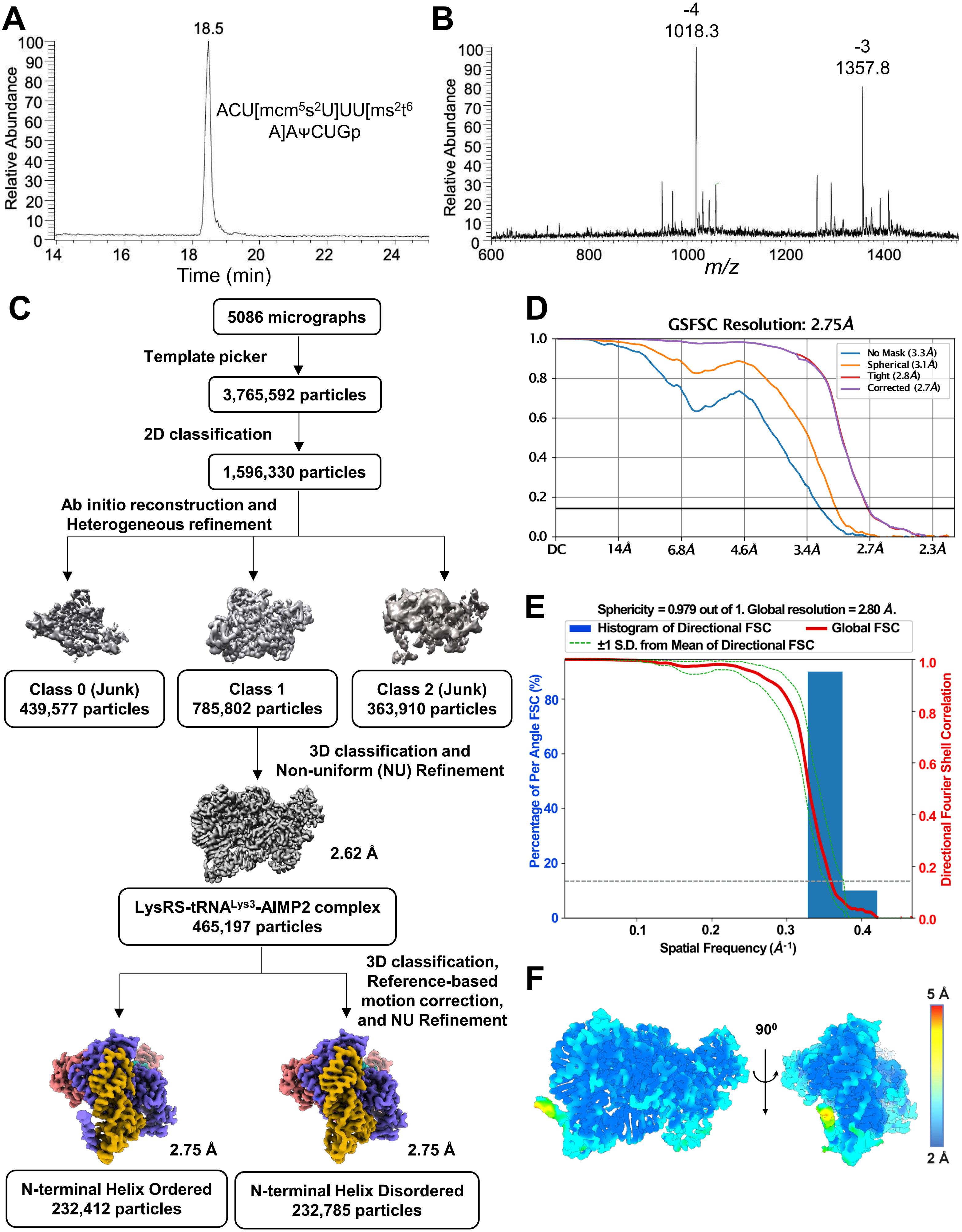

A
18.5
ACU[mcm5s2U]UU[ms2t6 A]AΨCUGp
Relative Abundance
Time (min)
B
-4
1018.3
-3
1357.8
Relative Abundance
m/z
C
5086 micrographs
Template picker
3,765,592 particles
2D classification
1,596,330 particles
Ab initio reconstruction and Heterogeneous refinement
Class 0 (Junk) 439,577 particles
Class 1 785,802 particles
Class 2 (Junk) 363,910 particles
3D classification and Non-uniform (NU) Refinement
2.62 Å
LysRS-tRNALys3-AIMP2 complex 465,197 particles
3D classification, Reference-based motion correction, and NU Refinement
2.75 Å
2.75 Å
N-terminal Helix Ordered 232,412 particles
N-terminal Helix Disordered 232,785 particles
D
GSFSC Resolution: 2.75Å
No Mask (3.3Å)
Spherical (3.1Å)
Tight (2.8Å)
Corrected (2.7Å)
DC
14Å
6.8Å
4.6Å
3.4Å
2.7Å
2.3Å
E
Sphericity = 0.979 out of 1. Global resolution = 2.80 Å.
Histogram of Directional FSC
Global FSC
±1 S.D. from Mean of Directional FSC
Percentage of Per Angle FSC (%)
Directional Fourier Shell Correlation
Spatial Frequency (Å-1)
F
90⁰
5 Å
2 Å

**Figure S1. (A)** Representative extracted ion chromatogram (EIC) of the summation of ions detected for the RNase T1 digestion product 'ACU[mcm$^5$s$^2$U]UU[ms$^2$t$^6$A]AYCUGp' from cellular modified tRNA$^{Lys3}$ eluting at 18.48 min. **(B)** Mass spectrum of the RNase T1 digestion product 'ACU[mcm$^5$s$^2$U]UU[ms$^2$t$^6$A]AYCUGp' from tRNA$^{Lys3}$. The calculated molecular weight for this digestion product is 4077.6 and the -3 and -4 charge states are labeled in this spectrum. **(C)** Overall data processing workflow for the LysRS-cellular modified tRNA$^{Lys3}$-AIMP2 cryo-EM reconstruction. **(D)** The gold-standard Fourier Shell Correlation (GS-FSC) curve for the LysRS-cellular modified tRNA$^{Lys3}$-AIMP2 reconstruction is shown. **(E)** 3D-FSC plot for the LysRS-cellular modified tRNA$^{Lys3}$-AIMP2 reconstruction showing the directional resolution anisotropy. An overall sphericity of 0.98 shows that the reported global resolution of 2.8 Å is uniform in all three dimensions and the sample does not show major preferred orientations. **(F)** The local resolution for the final cryo-EM reconstruction of the LysRS-cellular modified tRNA$^{Lys3}$-AIMP2 complex is shown in a color-coded format.

### Figure S2

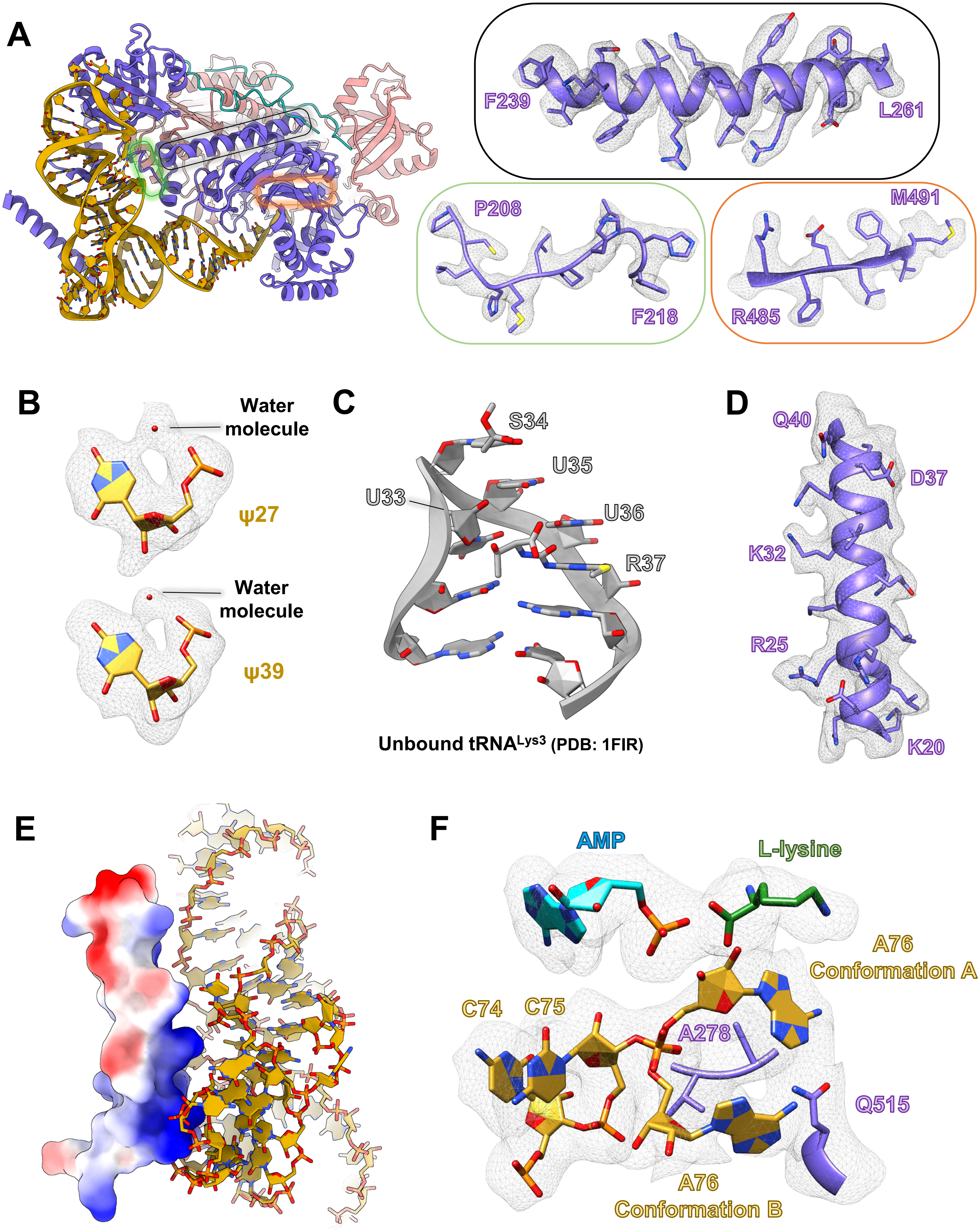

A
F239
L261
P208
F218
M491
R485
B
Water
molecule
ψ27
Water
molecule
ψ39
C
S34
U35
U33
U36
R37
Unbound tRNALys3 (PDB: 1FIR)
D
Q40
D37
K32
R25
K20
E
F
AMP
L-lysine
A76
Conformation A
C74
C75
A278
Q515
A76
Conformation B

**Figure S2. (A)** A few representative examples for the cryo-EM map and the corresponding modeled regions are shown. The location of the highlighted regions is indicated in the overall structural model using color coded boxes. Magnified views show the atomic model along with the corresponding cryo-EM map density in mesh. **(B)** Water molecule is often coordinated by the H1N1 imino group and the 5′-phosphate group of pseudouridine nucleotides, a feature not observed for uridine nucleotides. ψ27 and ψ39 are shown along with the coordinated water molecule. Corresponding cryo-EM map densities are shown in mesh. **(C)** The anticodon stem loop of cellular modified $tRNA^{Lys3}$ (PDB:1FIR) in the unbound form is shown highlighting the stacking interactions between the five anticodon loop bases (33-37). **(D)** The atomic model for the h-LysRS NTD is shown along with the corresponding cryo-EM map density in mesh. **(E)** The N-terminal subdomain of LysRS (15-43) is shown in surface view and color-coded by electrostatic potential (blue – basic; red – acidic; white – neutral). The corresponding interacting region of modified $tRNA^{Lys3}$ is shown in cartoon stick format. **(F)** The active site of the LysRS-cellular modified $tRNA^{Lys3}$-AIMP2 complex (docked state) is shown along with the corresponding cryo-EM map density to highlight the two alternate conformations of the A76 nucleotide of $tRNA^{Lys3}$.

### Figure S3

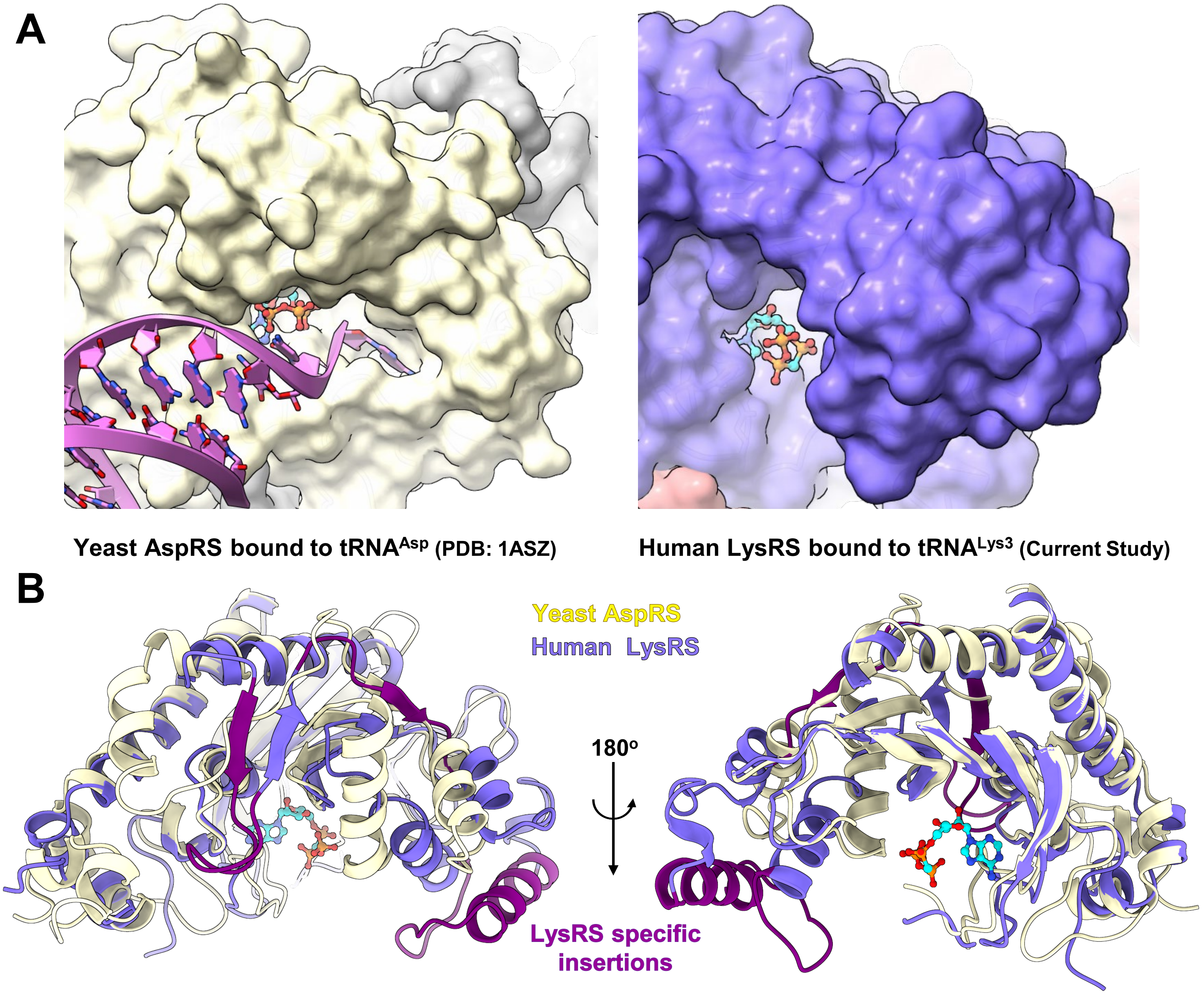


**C**

```
sp|Q15046|SYK_HUMAN      IETPMMNIIPGGAVAKPFITYHNELDMNLYMRIAPELYHKMLVVGGIDRVYEIGRQFRNE  325
sp|P04802|SYDC_YEAST     VHTPKLLGAPSEGGSSVFE--VTYFKGKAYLAQSPQFNKQQLIVADFERVYEIGPVFRAE  327
                         :.** :   *. . :. *    . :. : *:  :*:: :: *:*..::******  ** *

sp|Q15046|SYK_HUMAN      GIDL-THNPEFTTCEFYMAYAD-YHDLMEITEKMVSGMVKHITGSYKVTYHPDGPEGQAY  383
sp|P04802|SYDC_YEAST     NSNTHRHMTEFTGLDMEMAFEEHYHEVLDTLSELFVFIFSELPKRF------------AH  375
                         . :   *  ***  :: **: : **::::  .::.  :...:   :            *:

sp|Q15046|SYK_HUMAN      DVDFTPPFRRINMVEELEKALGMKLPETNLFETEETRKILDDICVAKAVECPPPRTTARL  443
sp|P04802|SYDC_YEAST     EIEL---------VRKQYPVEEFKLPKDGKMVRLTY---------KEGI------EMLRA  411
                         ::::         *.:   .  :***: . :               :.:         *
```

**Figure S3. (A)** A side-by-side view of the active sites of related class-II AARS, yeast AspRS (left panel, yellow) and human LysRS (right panel, purple), in the ATP bound state is shown. The entrance for the LysRS active site is considerably narrow compared to yeast AspRS, and unlike yeast AspRS, the simultaneous binding of ATP and the tRNA 3'-CCA end is incompatible in the LysRS active site. **(B)** The structures of the yeast AspRS-tRNA$^{Asp}$ complex (yellow) and human LysRS-tRNA$^{Lys3}$ complex (purple) were aligned and the aminoacylation domains (AAD) for both are shown in cartoon format. Despite high degrees of conservation, LysRS AAD has insertions (highlighted in deep purple) that lead to a narrow entrance to its active site, in contrast to AspRS. **(C)** Sequence alignment between yeast AspRS (Uniprot: P04802) and human LysRS (Uniprot: Q15046) was carried out using Clustal Omega and is shown to highlight the LysRS-specific insertions (highlighted in purple box).

### Figure S4

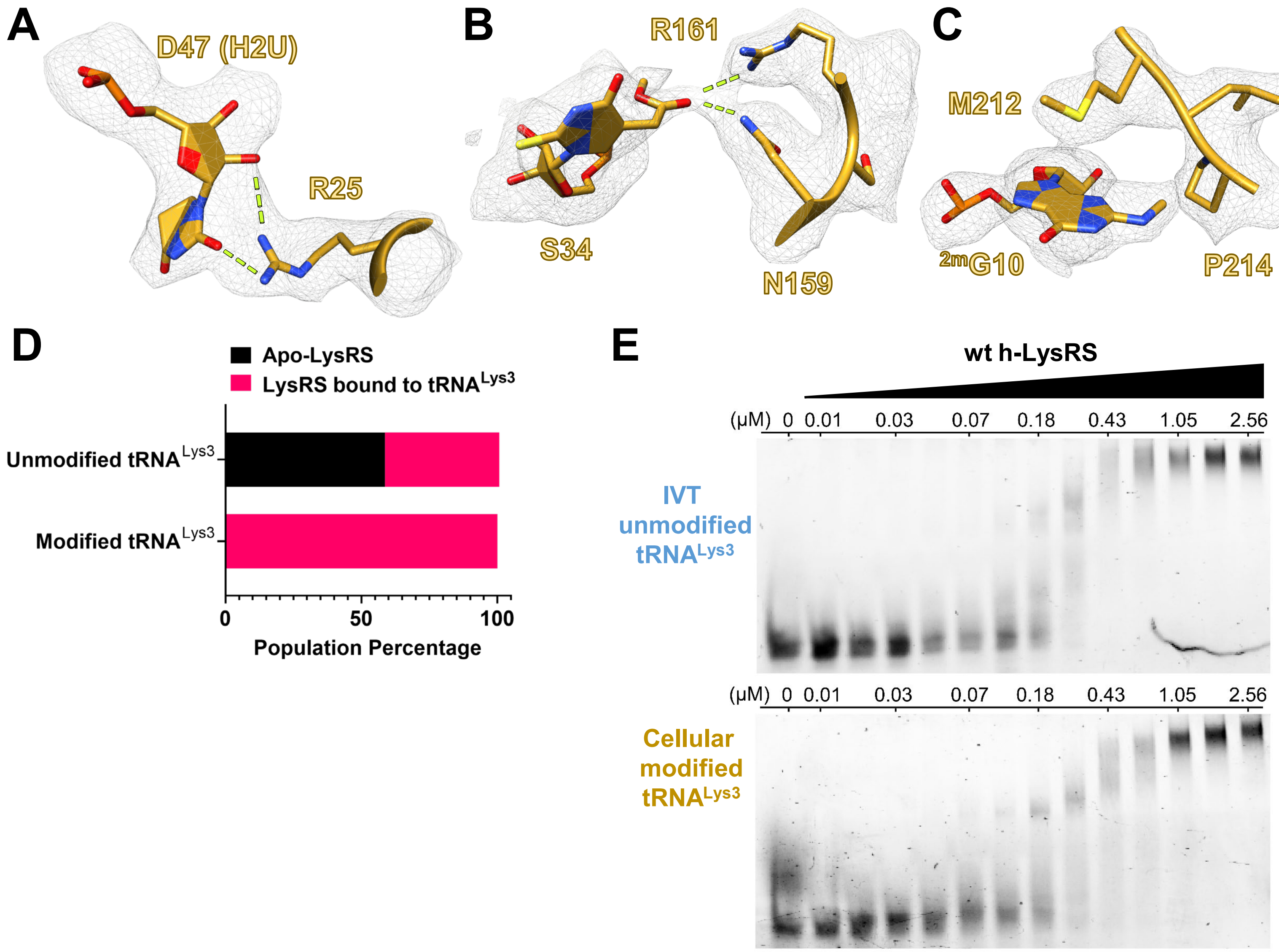


**Figure S4. (A-C)** LysRS specifically recognizes the tRNA$^{Lys3}$ modifications and a few representative examples are shown along with the corresponding cryo-EM map density. The three examples shown here are D47 (5,6 dihydrouracil) (A), S34 (5-methoxycarbonylmethyl-2-thiouridine) (B), and $^{m2}$G10 (2-methyl guanosine) (C) **(D)** Population statistics of h-LysRS from the cryo-EM dataset for cellular modified tRNA$^{Lys3}$ (+AIMP2) and IVT unmodified tRNA$^{Lys3}$ (+AIMP2) are shown in bar chart form. **(E)** A representative EMSA gel showing h-LysRS titration against IVT unmodified and cellular modified tRNA$^{Lys3}$.

### Figure S5

## A

```
sp|O43776|SYNC_HUMAN      MVLAELYVSDREGSDATGDGTKEKPFKTGLKALMTVGKEPFPTIYVDSQKENERWNVISK
sp|Q15046|SYK_HUMAN       -------MAAVQAAEVKVDGSEPKLSKNELKRRLKAEKKV-------AEKEAKQKELSEK
sp|P14868|SYDC_HUMAN      ----------------MPSASASRKSQEKPREIMDAAEDY----------AKERYGISSM
                                            ..:  .  :   .  : . :.             :.  : .

sp|O43776|SYNC_HUMAN      SQLKNIKKMWHREQMKSESREKKEAEDSLRREKNLEEAKKITIKNDPSLPEPKC--
sp|Q15046|SYK_HUMAN       QLSQATAAATNHTTDNGVGPEEESVDPNQYYKIRSQAIHQLKVNGEDPYPHKFHVD
sp|P14868|SYDC_HUMAN      IQSQEK-------------------------------------------PDRVL--
                             :                                             *
```

## B

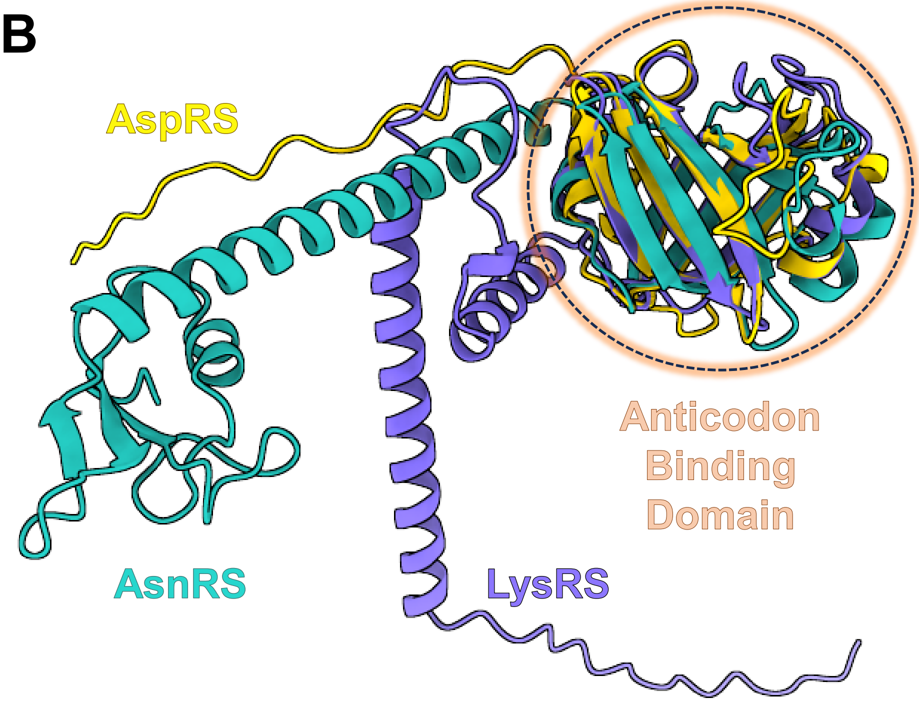


**Figure S5. (A)** The N-terminal extension domains of LysRS (Q15046, 1-102), AspRS (P14868, 1-45), and AsnRS (O43776, 1-114) were aligned using MUSCLE multiple sequence alignment algorithm. **(B)** The AlphaFold models for the N-terminal extension domains and anticodon-binding domains of LysRS (purple), AspRS (yellow), and AsnRS (sea green) were aligned in ChimeraX and are depicted in ribbon cartoon. The anticodon-binding domains of the three class-IIb AARS align well and are highlighted (orange circle).

### Table S1

#### Mass Spectrometry Analysis for Modified Cellular tRNA$^{Lys3}$

| Digestion Product | Theoretical *m/z* | Experimental *m/z* |
|---|---|---|
| **CCCGp** | (-1) 1277.8<br>(-2) 638.38 | (-1) 1277.2<br>(-2) 638.25 |
| **AUA[m$^2$G]p** | (-1) 1340.8<br>(-2) 669.91 | (-1) 1340.1<br>(-2) 669.83 |
| **CUCAGp** | (-1) 1608.0<br>(-2) 803.48 | (-1) 1607.0<br>(-2) 803.25 |
| **DCGp** | (-1) 975.59 | (-1) 975.17 |
| **DAGp** | (-1) 999.61 | (-1) 999.17 |
| **CAΨCAGp** | (-2) 968.08<br>(-3) 645.05 | (-2) 967.67<br>(-3) 645.00 |
| **ACU[mcm$^5$s$^2$U]UU[ms$^2$t$^6$A]AΨCUGp** | (-3) 1358.2<br>(-4) 1018.4 | (-3) 1357.8<br>(-4) 1018.3 |
| **[m$^7$G]UC[m$^5$C]AGp** | (-2) 990.11 | (-2) 990.67 |
| **[m$^5$Um]ΨCA[m$^1$A]Gp** | (-2) 989.62<br>(-3) 659.41 | (-2) 989.25<br>(-3) 659.33 |
| **UCCCUGp** | (-2) 944.55<br>(-3) 629.36 | (-2) 944.33<br>(-3) 629.33 |
| **UUCGp** | (-2) 639.36 | (-2) 639.33 |
| **CCA-OH** | (-1) 876.60 | (-1) 876.17 |

Table S1. All RNase T1 digestion products from cellular modified tRNA$^{Lys3}$ detected by LC-MS. The sequences were confirmed by MS/MS data, except for the anticodon and variable loop region, which were analyzed manually. Both theoretical *m/z* values and those experimentally determined are listed.

### Table S2

#### Cryo-EM Data Collection and Processing

| | h-LysRS bound to modified $tRNA^{Lys3}$ and AIMP2 | h-LysRS bound to IVT $tRNA^{Lys3}$ | h-LysRS bound to IVT $tRNA^{Lys3}$ | h-LysRS bound to IVT $tRNA^{Lys3}$ and AIMP2 |
|---|---|---|---|---|
| **Ligands** | AMP and L-lysine | AMPCPP and L-lysine | AMP and L-lysine | AMP and L-lysine |
| **Number of micrographs** | 4746 | 4643 | 5320 | 4045 |
| **Magnification** | 81,000x | 81,000x | 81,000x | 81,000x |
| **Voltage (kV)** | 300 kV | 300 kV | 300 kV | 300 kV |
| **Electron exposure ($e^{-}/Å^2$)** | 50 $e^{-}/Å^2$ | 50 $e^{-}/Å^2$ | 50 $e^{-}/Å^2$ | 50 $e^{-}/Å^2$ |
| **Defocus range (µm)** | - 1.0 to -2.0 | - 1.0 to -2.0 | - 1.0 to -2.0 | - 1.0 to -2.0 |
| **Pixel size (Å)** | 1.07 Å physical pixel size | 1.07 Å physical pixel size | 1.07 Å physical pixel size | 1.07 Å physical pixel size |
| **Microscope** | Titan Krios | Titan Krios | Titan Krios | Titan Krios |
| **Detector** | Gatan K3 | Gatan K3 | Gatan K3 | Gatan K3 |
| **Data Collection Software** | EPU | EPU | EPU | EPU |
| **Data Processing Software** | cryoSPARC v4 | cryoSPARC v4 | cryoSPARC v4 | cryoSPARC v4 |
| **Symmetry imposed** | C1 | C1 | C1 | C1 |
| **Initial particle images (no.)** | 785,802 | 881,907 | 1,211,344 | 601,206 |
| **Final particle images (no.)** | 232,412 | 430,648 | 302,069 (Docked state) 267,590 (Undocked state) | 165,298 |
| **FSC threshold and Map resolution (Å)** | 0.143 FSC cutoff 2.7 Å (Docked state) | 0.143 FSC cutoff 2.8 Å (Undocked state) | 0.143 FSC cutoff 3 Å (Docked state) 3.1 Å (Undocked state) | 0.143 FSC cutoff 4 Å (Docked state) |
| **Resolution range (Å)** | 2.1 Å – 6.5 Å | 2.3 Å – 10 Å | 2.3 Å – 10 Å | 2.4 Å – 11 Å |

Table S2. Details of cryo-EM data collection and data processing for the four different datasets are presented in a tabular format.

### Table S3

#### Model Building Statistics and Accession Codes

| | LysRS bound to cellular modified tRNA$^{Lys3}$ and AIMP2 (Docked State) | LysRS bound to IVT tRNA$^{Lys3}$ (Undocked State) | LysRS bound to IVT tRNA$^{Lys3}$ (Docked State) | LysRS bound to IVT tRNA$^{Lys3}$ (Undocked State) |
|---|---|---|---|---|
| **Ligands** | AMP and L-lysine | AMPCPP and L-lysine | AMP and L-lysine | AMP and L-lysine |
| **Deposition ID** | PDB: 9DPL<br>EMD-47106 | PDB: 9DPB<br>EMD-47101 | PDB: 9DPA<br>EMD-47100 | PDB: 9DOW<br>EMD-47094 |
| **Map sharpening B factor (Å²)** | - 55 | - 76 | - 45 | - 33 |
| **Initial Model Used** | PDB: 6ILD, 1FIR | PDB: 6ILD, 1FIR | PDB: 6ILD, 1FIR | PDB: 6ILD, 1FIR |
| **Model composition** | | | | |
| **Non-hydrogen atoms** | 10555 | 9836 | 9878 | 9816 |
| **Protein residues** | 1092 | 1013 | 1013 | 1013 |
| **Nucleotides** | 76 | 73 | 76 | 73 |
| **Metal Ions** | 4 | 5 | 1 | 1 |
| **R.m.s. deviations** | | | | |
| **Bond lengths (Å)** | 0.01 | 0.006 | 0.007 | 0.005 |
| **Bond angles (°)** | 1.327 | 1.002 | 1.032 | 0.955 |
| **Validation** | | | | |
| **MolProbity score** | 1.68 | 1.85 | 1.93 | 1.89 |
| **Clashscore** | 9.12 | 9.93 | 10.63 | 10.10 |
| **Ramachandran plot** | | | | |
| **Favored (%)** | 96.94 | 96.72 | 96.43 | 96.62 |
| **Allowed (%)** | 3.06 | 3.28 | 3.57 | 3.38 |
| **Disallowed (%)** | 0.00 | 0.00 | 0.00 | 0.00 |

Table S3. Model building, data deposition and validation statistics.

### Table S4

#### Sequence Information and Coverage for the Structural Models

| Name | Alternative Name | Chain ID | Length (residues) | Modeled residues | UniProt ID (proteins) |
|---|---|---|---|---|---|
| **h-LysRS bound to cellular modified tRNA$^{Lys3}$ and AIMP2-N36 (AMP and L-lysine) PDB: 9DPL** | | | | | |
| **h-LysRS** | **KARS** | A | 597 | 14-43, 69-579 | Q15046 |
| **h-LysRS** | **KARS** | B | 597 | 71-577 | Q15046 |
| **tRNA$^{Lys3}$** | **-** | C | 76 | 76 | HG983931.1 (GenBank) |
| **AIMP2-N36** | **-** | D and E | 36 | 1-21 | Q13155 |
| **h-LysRS bound to IVT tRNA$^{Lys3}$ and AIMP2 (AMPCPP and L-lysine) [Undocked State] PDB: 9DPB** | | | | | |
| **h-LysRS** | **KARS** | A and B | 597 | 72-577 (A) and 72-576 (B) | Q15046 |
| **tRNA$^{Lys3}$** | **-** | C | 76 | 73 | HG983931.1 (GenBank) |
| **h-LysRS bound to IVT tRNA$^{Lys3}$ and AIMP2 (AMP and L-lysine) [Docked State] PDB: 9DPA** | | | | | |
| **h-LysRS** | **KARS** | A and B | 597 | 72-576 (A, B) | Q15046 |
| **tRNA$^{Lys3}$** | **-** | C | 76 | 76 | HG983931.1 (GenBank) |
| **h-LysRS bound to IVT tRNA$^{Lys3}$ and AIMP2 (AMP and L-lysine) [Undocked State] PDB: 9DOW** | | | | | |
| **h-LysRS** | **KARS** | A and B | 597 | 72-576 (A) and 72-577 (B) | Q15046 |
| **tRNA$^{Lys3}$** | **-** | C | 76 | 73 | HG983931.1 (GenBank) |

Table S4. Details of the four structural models including the Chain IDs, total length, modeled residues, and Uniprot/Genbank IDs are listed.

### Table S5

#### Conformational States and Population Statistics

| Dataset | Ligand | Apo-LysRS | h-LysRS-tRNA$^{Lys3}$ complex | |
|---|---|---|---|---|
| | | | Undocked State | Docked State |
| **h-LysRS with cellular modified tRNA$^{Lys3}$ and AIMP2** | AMP and L-lysine | 0 | 0 | 465,197 (100%) |
| **h-LysRS with IVT tRNA$^{Lys3}$** | AMPCPP and L-lysine | 458,550 (52%) | 430,648 (48%) | 0 |
| **h-LysRS with IVT tRNA$^{Lys3}$** | AMP and L-lysine | 268,855 (32%) | 267,590 (32%) | 302,069 (36%) |
| **h-LysRS with IVT tRNA$^{Lys3}$ and AIMP2** | AMP and L-lysine | 235,233 (59%) | 0 | 165,298 (41%) |

Table S5. Details of the distinct conformational states resolved in the four cryo-EM datasets. The final number of particles for each of the indicated conformational states are tabulated. The relative percentage of the particle stacks belonging to each conformational state in a dataset is presented in parenthesis.

**Movie S1. h-LysRS remodels the anticodon loop of tRNA$^{Lys3}$.** This video highlights the remodeling of modified tRNA$^{Lys3}$ upon binding by h-LysRS. All 5 nucleotides in the anticodon loop are unstacked and involved in binding h-LysRS. The video was made using UCSF ChimeraX.

**Movie S2. tRNA$^{Lys3}$ modifications mediate recognition by h-LysRS.** This video highlights the multiple identity elements used by h-LysRS for binding cellular modified tRNA$^{Lys3}$ and the integral role played by tRNA$^{Lys3}$ modifications. The video was made using UCSF ChimeraX.

**Movie S3. Conformational Landscape of tRNA$^{Lys3}$ aminoacylation.** This video highlights the 3'-CCA undocked state and 3'-CCA docked state conformations of h-LysRS-tRNA$^{Lys3}$ complex and how the ATP state in the active site governs the conformational dynamics. The ordering of the two catalytic steps in tRNA$^{Lys}$ aminoacylation by h-LysRS is markedly different to previously reported aminoacylation pathway of yeast AspRS, a related class-II aminoacyl-tRNA synthetase. The video was made using UCSF ChimeraX.

**Movie S4. Docked state of h-LysRS complex with cellular modified tRNA$^{Lys3}$.** This video highlights the 3'-CCA docked state conformation of the h-LysRS-modified tRNA$^{Lys3}$ complex. A76 base is flipped placing the 3'-hydroxyl in close proximity to AMP (cyan) and L-lysine (green). The video was made using UCSF ChimeraX.
